# Population differentiation of polygenic score predictions under stabilizing selection

**DOI:** 10.1101/2021.09.10.459833

**Authors:** Sivan Yair, Graham Coop

## Abstract

Given the many small-effect loci uncovered by genome-wide association studies (GWAS), polygenic scores have become central to the drive for genomic medicine and have spread into various areas including evolutionary studies of adaptation. While promising, these scores are fraught with issues of portability across populations, due to mis-estimated effect sizes and missing causal loci across populations unrepresented in large-scale GWAS. The poor portability of polygenic scores at first seems at odds with the view that much of common genetic variation is shared among populations. Here we investigate one potential cause of this discrepancy, stabilizing selection on complex traits. Somewhat counter-intuitively, while stabilizing selection to the same optimum phenotype leads to lower phenotypic differentiation among populations, it increases genetic differentiation at GWAS loci because it accelerates the turnover of polymorphisms underlying trait variation within populations. We develop theory to show how stabilizing selection impacts the utility of polygenic scores when applied to unrepresented populations. Specifically, we quantify their reduced prediction accuracy and find they can substantially overstate average genetic differences of phenotypes among populations. Our work emphasizes stabilizing selection to the same optimum as a useful null evolutionary model to draw connections between patterns of allele frequency and polygenic score differentiation.

## 2 Introduction

Lewontin’s foundational early survey of human genetic variation found that much of genetic variation is found within human populations rather than between human populations, i.e. that human populations are only weakly differentiated at the level of individual loci and most common variation is shared among populations (Lewontin, 1972). Thus it strongly refuted an alternative view that genetic variation would be partitioned among mostly invariable populations and has become a classic work discrediting discrete human races. Lewontin’s finding is reflected by low estimates of *F*_*ST*_ among populations (Li *et al*., 2008), i.e. only a small proportion (∼ 10%) of the total allelic variance is attributable to differences in frequency between populations. Lewontin’s early work relied on breakthroughs in measures of protein variation via electrophoretic mobility and earlier surveys of blood groups. Since then, the findings of low *F*_*ST*_ have been found to hold over many different markers across the genome (Barbujani *et al*., 1997; Jorde *et al*., 2000; Rosenberg *et al*., 2002; Conrad *et al*., 2006; Li *et al*., 2008; Bergström *et al*., 2020). While some loci that are highly differentiated among human populations have been uncovered (e.g. underlying loci in skin pigmentation and infectious disease immunity; see Fan *et al*., 2016, for a review), there are relatively few such strongly selected loci (Coop *et al*., 2009; Hernandez *et al*., 2011).

Lewontin’s results also have important implications for our *a priori* expectations of the partitioning of phenotypic variation within and among human populations. That is because for phenotypes evolving neutrally in diploids, we expect the proportion of additive genetic variance attributable to among-population differences to be ≈ 2*F*_*ST*_ when all individuals are measured in a common set of environments (Wright, 1951; Lande, 1976; Rogers and Harpending, 1983; Lande, 1992; Whitlock, 1999; Edge and Rosenberg, 2015). Thus in humans, we expect only ∼ 18% of additive genetic variance to be due to among population differences when measured in a common set of environments. Compared to this null, phenotypes subject to divergent selection among populations can be over-dispersed, while phenotypes subject to stabilizing selection with the same selective optimum are expected to be less differentiated among populations. To distinguish among potential contributors to genetic differences among populations, researchers often turn to settings like common gardens in an effort to eliminate among-population environmental variation. However, it is obviously not feasible to measure human phenotypes in a common environment (a major drawback of studies that investigated population level phenotypic variation; Relethford and Lees, 1982; Chakraborty, 1990). Thus for the vast majority of complex traits, we do not know the role of genetics, let alone natural selection, in explaining phenotypic differences among human populations.

These questions have received renewed interest in human genetics due to genome-wide association studies (GWAS), which have found common variation associated with many phenotypes within populations. GWAS have revealed that most phenotypes are highly polygenic within populations (Loh *et al*., 2015; Shi *et al*., 2016; Boyle *et al*., 2017) and confirmed that much of the genetic variance is additive (reviewed in Hill *et al*., 2008). These observations have motivated phenotype prediction through additive genetic values, the additive contribution of polymorphisms to phenotypic differences among individuals. One common approach to predict an individual’s additive genetic value is based on a polygenic score, the sum across trait-associated loci of genotypes weighted by their estimated effects. Predictions based on polygenic scores are being explored in a number of clinical settings and more generally as a tool for understanding the genetic basis of disease and phenotypic variability. However, the generalizability of polygenic scores across populations is a key concern because GWAS samples are strongly biased towards European populations and studies of other populations are much smaller (Li and Keating, 2014; Popejoy and Fullerton, 2016; Macarthur *et al*., 2017; Martin *et al*., 2019). There is wide agreement that these portability issues must be addressed if the future clinical use of polygenic scores is not to just further compound inequalities in healthcare (Li and Keating, 2014; Popejoy and Fullerton, 2016; Martin *et al*., 2019).

Currently, polygenic scores tend to poorly predict additive genetic values underlying traits, and therefore phenotypes, even in samples closely related to the GWAS population, because they aggregate across many loci with slightly mis-estimated effects. These problems increase as we move to populations that are genetically and environmentally more distant to the GWAS populations. For one, the associated loci are usually not the causal loci underlying trait variation; instead they tag the effects of linked causal sites. The interpretation of the effect size of an associated variant can be tricky because of: (i) linkage disequilibrium (LD), whereby it absorbs the effects at correlated causal sites (Vilhjálmsson *et al*., 2015; Horikoshi *et al*., 2017; Lam *et al*., 2019; Wang *et al*., 2020; Weissbrod *et al*., 2021); (ii) population stratification, whereby it absorbs the effects of covarying environments (Haworth *et al*., 2019; Kerminen *et al*., 2019; Sakaue *et al*., 2020; Trochet and Hussin, 2020; Isshiki *et al*., 2021); and (iii) gene-by-environment (GxE) or gene-by-gene (GxG) interactions, whereby its estimate is averaged over the interacting environmental contexts or genetic backgrounds in the sample (Brown *et al*., 2016; Horikoshi *et al*., 2017; Coram *et al*., 2017; Adhikari *et al*., 2019; Bentley *et al*., 2019; Galinsky *et al*., 2019; Grinde *et al*., 2019; Veturi *et al*., 2019; Wojcik *et al*., 2019; Mostafavi *et al*., 2020; Mathieson, 2021; Patel *et al*., 2021). As all of these factors can and will vary across populations, the effect sizes of alleles will differ among them and so polygenic scores will have lower prediction accuracy, i.e. imperfect portability, across populations.

Even with perfectly estimated effects at the causal loci with significant trait associations, the prediction accuracy of polygenic scores will be limited because GWAS only identify loci with common alleles that contribute enough variance to exceed some significance threshold determined by the sample size. Therefore, an allele that is rare in the GWAS sample but common elsewhere will not be discovered, leading to a greater reduction in the variance accounted for, or prediction accuracy, in unrepresented populations (Martin *et al*., 2017a,b; Curtis, 2018; Kim *et al*., 2018; Bentley *et al*., 2019; Wojcik *et al*., 2019; Wang *et al*., 2020; Conti *et al*., 2021). Indeed, many variants contributing to trait variation in European GWAS samples are not at a high enough frequency to be detected in other populations, suggesting different sets of polymorphisms contribute to the trait variance in different populations (Liu *et al*., 2015; Durvasula and Lohmueller, 2021). Genetic differentiation likely contributes to the reduction in the prediction accuracy of polygenic scores in unrepresented populations, as groups with increasing genetic distance from GWAS samples experience a greater loss in prediction accuracy (Scutari *et al*., 2016; Bitarello and Mathieson, 2020; Cavazos and Witte, 2021; Privé *et al*., 2022).

The factors that reduce the utility of polygenic scores for individual level prediction may also complicate the interpretation of average polygenic score differences across populations. Such issues arise in studies of adaptation that use polygenic scores to assess the contribution of selection to the genetic basis of phenotypic differentiation among human populations. An early application of this approach found polygenic signals of selection on height within Europe (Turchin *et al*., 2012; Berg and Coop, 2014), where polygenic scores were over-differentiated among populations compared to the neutral prediction based on *F*_*ST*_. Importantly, the null distribution of this test of neutrality at the level of the phenotype does not rely on accurate polygenic scores and so changes in LD, GxG, and GxE should not cause false signals. However, the results are very sensitive to slight biases in estimated effect sizes due to population structure, and indeed the signal of polygenic selection on height turned out to be almost entirely due to stratification (Berg *et al*., 2019; Sohail *et al*., 2019; Refoyo-Martínez *et al*., 2020). More generally, the imperfect portability of polygenic scores, and the fact that the mean environmental contribution to phenotypes can vary greatly between populations, raises significant concerns about over-interpreting differences in mean polygenic scores as genetic differences in the average phenotype among populations (Berg and Coop, 2014; Novembre and Barton, 2018; Coop, 2019; Rosenberg *et al*., 2019; Harpak and Przeworski, 2021).

The limited generalizability of GWAS results for phenotype prediction and comparison beyond the study sample may appear to contradict Lewontin’s observation of minor allele frequency differences between populations. However, allele frequency differentiation for complex traits under selection can occur at a rate faster than drift. This would lead to noisy estimates of additive genetic values, and so reduced prediction accuracy of polygenic scores, for unrepresented populations. In order to understand the impact of this turnover on portability, we need models of allele frequency differentiation that are informed by plausible forms of natural selection on complex traits.

Stabilizing selection with a constant fitness optimum is a sensible null model for the evolution of complex traits. Indeed, studies of its influence on allelic dynamics have set a conceptual foundation for interpreting and designing GWAS within populations (Simons *et al*., 2018). Under stabilizing selection, intermediate trait values have the highest fitness, with decreasing fitness with distance from that optimum (Figure 1A). Many quantitative traits have been shown to experience stabilizing selection in humans (e.g. Sanjak *et al*., 2018) as well as across many other species (e.g. Bumpus, 1899; Kingsolver *et al*., 2001; de Villemereuil *et al*., 2020). Stabilizing selection also contributes to the lack of variation in morphological traits within and between closely related species and the morphological constancy of traits in the fossil record (Gingerich, 1983; Weber, 1990; Barton and Keightley, 2002; Hill and Kirkpatrick, 2010; Houle *et al*., 2017).

**Figure 1:**
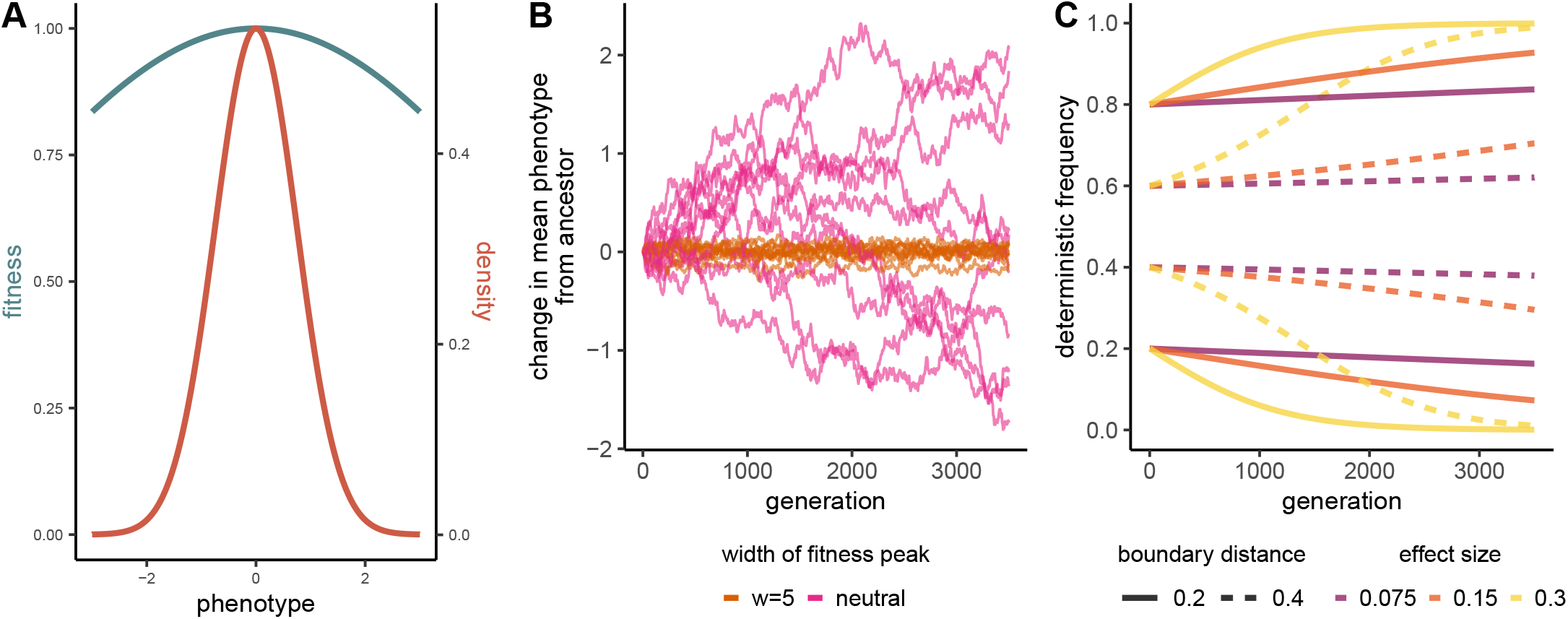
**A)** Gaussian fitness function for phenotypic stabilizing selection (teal) and resulting density of simulated phenotypes in the population (coral). Lines correspond to the intermediate strength of stabilizing selection that we simulated (*w* = 5, where *w* determines the width of the fitness peak, see Section 3). Note how stabilizing selection keeps the genetic variance of the population (*V*_*P*_) small compared to the width of the selection peak (*w*). **B)** Change in mean phenotype from the ancestor (generation 0) under stabilizing selection (orange) and neutral phenotypic evolution (pink). Each line corresponds to a single simulation; here 10 simulations are shown for each strength of selection. **C)** Deterministic allele frequency trajectories, assuming an infinite population size, based on the underdominant model, at a locus that contributes to the variance of a trait under stabilizing selection (here *w* = 5). The trajectory differs according to effect size and starting frequency. Note the symmetry for starting frequencies that are the same distance to their closest boundaries.

Here, we show how stabilizing selection on complex traits reduces portability and increases the chance of false signals of directional polygenic adaptation, despite low overall genetic differentiation and no genetically-based trait differentiation among populations. Under parameter ranges estimated from empirical studies, we combine simulations of stabilizing selection on complex traits with analytical models of its effect on genetic differentiation to investigate how stabilizing selection drives these results. We focus only on scenarios in which the trait optimum is shared among populations, leading to levels of trait differentiation among populations much smaller than neutrality. In our baseline scenario, stabilizing selection occurs on a single additive trait, where alleles have the same effect in different populations (reflecting no GxG or GxE interactions and no pleiotropy, we relax these assumptions later).

We study differentiation between a pair of populations, either an ancestral and descendant population or a pair of contemporary populations, in which the results of a GWAS in one of those populations is used to make trait predictions in the other. In doing so we provide a polygenic score perspective on earlier investigations into the relationship between population structure, stabilizing selection, and quantitative trait variation (e.g. Cohan, 1984; Narain and Chakraborty, 1987; Lande, 1991; Goldstein and Holsinger, 1992; Latta, 1998; Le Corre and Kremer, 2003). We show how these factors reduce the prediction accuracy of polygenic scores and can readily lead to patterns of polygenic score differentiation rife with the potential for misinterpretation.

## 3 Model Background

An individual’s additive genetic value *G*_*i*_ is the sum of the additive effects of all alleles they carry,

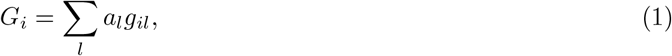

where *a*_*l*_ represents the additive effect of an allele relative to another at locus *l* and *g*_*l*_ is the number of copies of that allele carried by the individual. All of these loci denoted by *l* are polymorphic within a specified population from which the individual was drawn. Note that the additive genetic value does not represent an absolute measure of an individual’s phenotype, and instead represents the additive contribution of the polymorphisms they carry to their deviation from their population’s mean.

Under a constant selective environment, stabilizing selection keeps the population mean phenotype close to the optimum and decreases the phenotypic variance in the population because individuals on both tails of the distribution have lower fitness (Figure 1A,B; Wright, 1935; Lande, 1976; Lande and Arnold, 1983). To understand the process by which stabilizing selection reduces the phenotypic variance, we focus on the additive genetic variance,

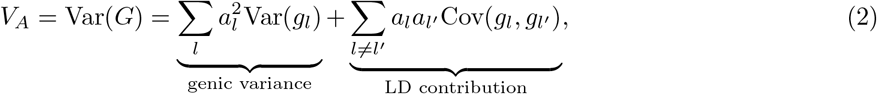

in which the first component refers to the additive *genic* variance (*V*_*a*_) and the second component accounts for the contribution of linkage disequilibrium (LD) among loci that contribute to the variance. In the short term, stabilizing selection reduces the phenotypic variance by generating negative LD between like-effect alleles, thereby limiting extreme phenotypes, which is known as the Bulmer effect (Bulmer, 1971). The additive genetic variance quickly reaches an equilibrium reflecting a balance between selection producing negative LD and recombination and chromosome segregation breaking up that generated LD (Turelli and Barton, 1994).

The long-term genetic response to stabilizing selection is driven by a reduction in the additive genic variance, in particular the variance in genotypes at a locus (Var(*g*_*l*_); the expected heterozygosity). Yet if the genetic basis of trait variation was truly infinitesimal, i.e. made up of loci of infinitely small effect, only genetic drift would erode variation in the long term. However, while the loci we discover with GWAS contribute very small effects, these effects are not infinitesimally small and so they can be directly acted on by selection (Simons *et al*., 2018; Sella and Barton, 2019). Selection at the individual loci underlying variation in the trait is in many cases well approximated by underdominant selection, in which the more common allele fixes and the minor allele is lost (Figure 2A; Wright, 1935; Robertson, 1956). Due to underdominant selection at the locus level, stabilizing selection removes polymorphisms at a faster rate than neutrality with selection coefficients proportional to their squared effect sizes (see Simons *et al*., 2018; Sella and Barton, 2019; Hayward and Sella, 2021, for recent applications of such models to understand GWAS variation within populations). Meanwhile, under moderate strengths of stabilizing selection and a constant environment, the population mean phenotype stays very close to the optimum through rapid, small fluctuations at many individual loci.

**Figure 2:**
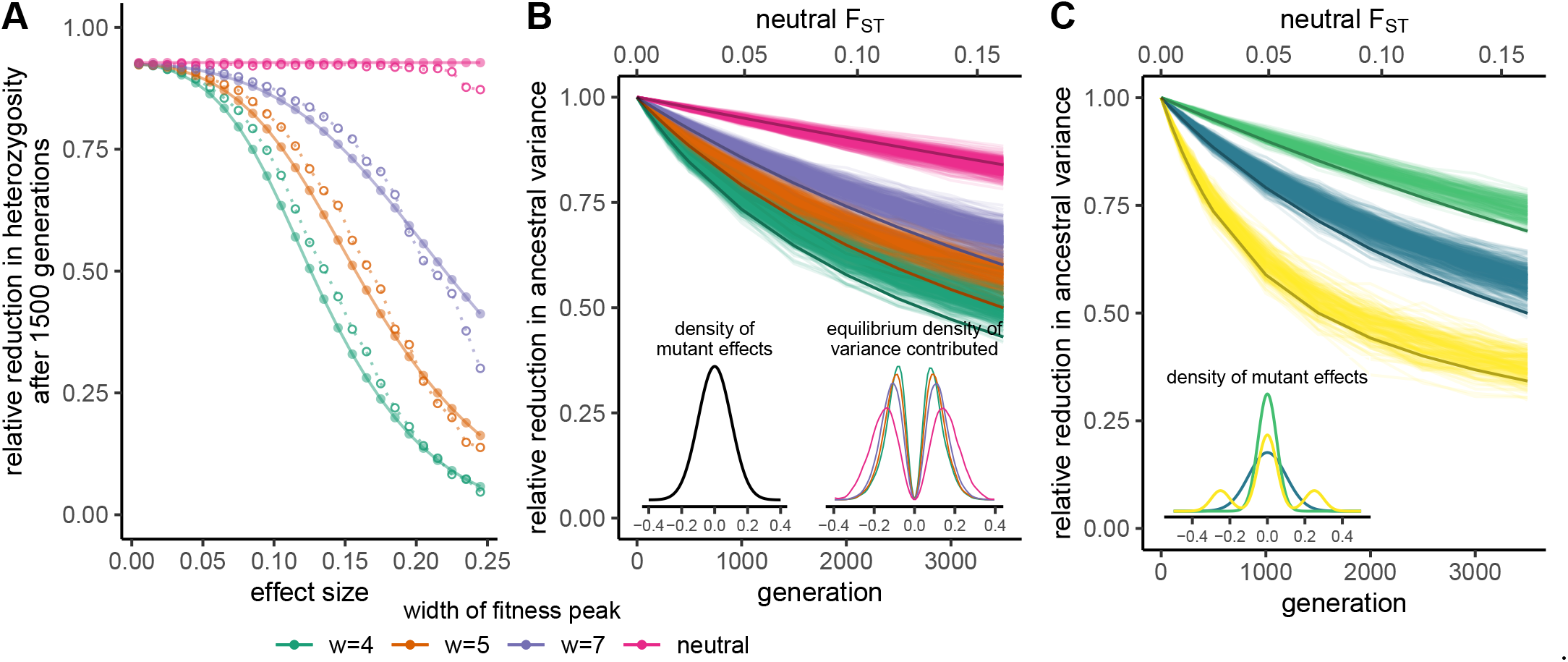
**A)** Reduction in heterozygosity at loci that contributed to the variance 1500 generations ago in a population with *N*_*e*_ = 10, 000. Each open point represents the midpoint of the effect size bin of width 0.01, within which we averaged heterozygosity from 200 simulations. Each filled point represents our analytical predictions for that midpoint. **B)** Reduction over time in the total variance contributed by polymorphisms in an ancestral population. Lighter lines show results from 200 simulations and darker lines show analytical predictions. The axis at the top shows the expected neutral *F*_*ST*_ between the ancestral and descendant population for the time scales of divergence shown on the bottom axis. The inset on the left shows the distribution from which mutant effect sizes are drawn. For the results shown we used a normal distribution with standard deviation of 0.1. The inset on the right shows the equilibrium density of variance contributed from each effect size. As the width of the fitness peak increases, and thus stabilizing selection weakens, mutations with large effects can drift to higher frequencies and contribute a greater proportion of the trait’s variance. **C)** The reduction over time in the total variance contributed by polymorphisms in an ancestral population for three different mutant effect size distributions, holding the strength of phenotypic stabilizing selection constant (*w* = 5). The inset shows the three mutant effect size distributions: two normal (green and blue) and a mixture of normals to produce a heavy-tailed distribution (yellow). The blue lines correspond to the same distribution used to produce sub-figure B, thus replicating the orange lines in sub-figure B. Lighter lines show results from simulations (100 each for yellow and green) and darker lines show analytical predictions. Predictions in **B** and **C** are based on the diffusion approximation, see Equations A.17 and A.18.

We note that the full analysis of models of stabilizing selection is challenging because changes in allele frequencies and covariances must be tracked over many loci (e.g. Turelli and Barton, 1990, 1994). Indeed, when we simulate physically linked loci with low recombination, we find that selection on an allele is weaker than we predict due to the persistence of selection-generated LD between alleles with opposing effects (the Bulmer effect; Figure S1). To understand the long-term turnover in the genetic basis of trait variation, from here on we focus on the additive genic variance, *V*_*a*_, under the assumption that loci contributing to the variance are physically unlinked.

We begin by thinking of an ancestral population at equilibrium and the loss of ancestral phenotypic variance over time. When the population is at mutation-drift-selection equilibrium, stabilizing selection and genetic drift remove variation such that as time goes on less and less of the variance in the descendant population is contributed by ancestral polymorphisms. Instead, the variance contributed by new mutations that are private to the descendant population will increase, replenishing what was removed by drift and selection.

We assume a model of Gaussian stabilizing selection where the width of the fitness peak is determined by *w* (*w*^2^ is equivalent to *V*_*S*_ in other models of stabilizing selection, e.g. Turelli, 1984), which will approximate any symmetric quadratic stabilizing selection model when the mean trait is close to its optimum. We also assume thousands of unlinked loci contributing to trait variation and explore dynamics resulting from three different mutant effect size distributions (Figure 2C); see Appendix A.1 for simulation details.

We can predict the reduction in additive genic variance contributed by ancestral variants (anc) in the descendant population (desc) after *t* generations for alleles with effect size *a* using a common approximation for the per generation loss (e.g. Keightley and Hill, 1988),

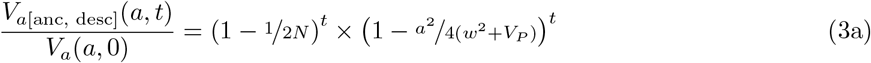

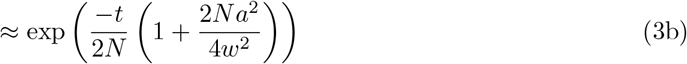

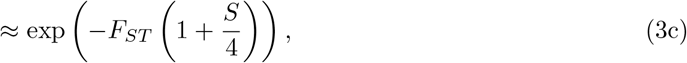

where *V*_*P*_ is the total population trait variance and *V*_*P*_ ≪ *w*^2^, *S* = 2*Na*^2^ */w*^2^ is the population-scaled selection coefficient of the allele, where *F*_*ST*_ ≈ *t/*2*N* for neutral polymorphisms. On the right-hand side of Equation 3a, the first bracketed term is the per generation reduction due to drift and the second term is the reduction due to stabilizing selection. Looking at the exponent in Equation 3c, we see that the decay of the variance (heterozygosity) contributed by alleles depends on *F*_*ST*_, but will be increased for alleles whose effect sizes are large enough that their population scaled selection is appreciable (*S >* 1). The above equation offers good intuition, however in the remainder of the main text we show results from a diffusion approximation that we developed that is better our purposes (extending from Simons *et al*., 2018, see Appendix A.3).

The reduction in additive genic variance contributed by a particular polymorphism with effect size *a* is the same as the reduction in heterozygosity at that site. Selection causes a stronger reduction when the fitness peak is narrower (i.e. when *w* is smaller) and when the allele’s effect is larger (Figure 2A; see Figure S2 for the approximation in Equation 3a). We can average the reduction in heterozygosity across all sites, weighting by the distribution of effect sizes and genic variance contributed by a given effect size, to predict the total remaining variance and thus total reduction in *V*_*a*_ over time (Figure 2B; Equation A.17). In the example shown in Figure 2, this total reduction tends to be weaker than what we see for the largest effect polymorphisms, because most sites that contribute to the variance are of small effect. This is because (i) under our mutational distribution most alleles have small effects, and (ii) under stabilizing selection, the equilibrium distribution of observed effects is more narrow than the distribution of mutant effects.

The form of the decay in ancestral variance strongly depends on the distribution of mutant effects. Consider a case in which a higher proportion of introduced mutations are strongly selected due to their large effects (*S >* 10). Some of these alleles are still capable of drifting to intermediate frequencies, and so a higher proportion of the ancestral variance will be due to larger effect polymorphisms (Simons *et al*., 2018). Therefore stabilizing selection will cause a steeper reduction in the variance they explain (Figure 2C). Moreover, if most of the other mutations are nearly neutral, the early and steep decline in ancestral variance will be followed by a decline more consistent with neutrality (yellow lines in Figure 2C; see Simons *et al*., 2018, for a discussion of selection regimes). The extent to which ancestral variation is removed determines the amount of shared additive genetic variance between diverging populations and thus the portability of polygenic scores.

## 4 Accuracy of polygenic score predictions

To understand the effect of stabilizing selection and drift on the prediction accuracy of polygenic scores in isolation from other sources of bias, we make the simplifying assumption that GWAS identify associations between polymorphisms and trait variation only at causal loci. For a GWAS within a population to identify a locus as being associated with the trait, the locus has to be polymorphic in that population and its phenotypic association has to achieve some level of statistical significance (i.e. contribute above some level of variance to the trait). For those causal loci with significant associations, we also assume their effects are estimated perfectly and for the moment that these true effects do not vary within the sample, population the sample was drawn from, or across populations (i.e. populations experience the same set of environments). We also assume their effects are strictly additive. At the trait-associated loci, one can sum the additive effects of all alleles an individual carries at a predefined set of markers to form a polygenic score.

We are interested in the reduction in prediction accuracy for a population not represented in the GWAS sample, relative to the prediction accuracy of those represented populations. To explore this we consider the genetic differentiation between a pair of populations, A and B, in which the GWAS sample is drawn from population A but not population B. When the effects of alleles do not vary between populations, we can quantify the reduction in phenotype prediction accuracy (*r*^2^) of polygenic scores, compared to using additive genetic values, constructed for any population as

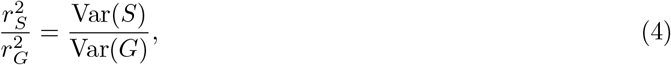

where *S*_*i*_ is an individual’s polygenic score and *G*_*i*_ is their additive genetic value. Since an individual’s polygenic score is only part of their additive genetic value when a GWAS estimates true effects, this reduction can be understood as the proportion of the total additive genetic variance, or proportion of the heritability, explained by GWAS-significant sites (Appendix A.2). The ratio of 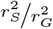 in population B to population A quantifies the reduction in prediction accuracy due to a lack of GWAS representation for population B. If these populations experience the same selective environment, we expect the same Var(*G*) for each population, so this reduction due to a lack of representation would simply be the ratio of the variances explained by polygenic scores.

Note that our definition of the prediction accuracy in population B is the squared correlation of the deviation of an individual’s polygenic score with its deviation in phenotype, where both of these deviations are with respect to the mean phenotype in population B. This definition matches typical polygenic score practices where predictions are statements about the departure of an individual from their ancestry group’s mean genetic value, rather than a prediction of their departure from the mean genetic value of the GWAS population. However, this definition of prediction accuracy does not include discrepancies from the evolution of mean polygenic scores and phenotypes among populations, meaning it does not account for a systematic shift in polygenic scores between populations. We turn to this point in Section 5.

### 4.1 Ascertainment of all causal loci in GWAS sample

We begin with the simplified case in which all causal loci that are polymorphic in population A have been identified, such that polygenic scores equal the additive genetic values in population A. As polygenic scores from ancient DNA are being used to investigate the phenotypic diversity of ancient human populations (Irving-Pease *et al*., 2021), we first consider the prediction accuracy in a population ancestral to population A of a polygenic score constructed using variants found in population A. The reduction in polygenic score prediction accuracy for this ancestral population is approximately the reduction in variance contributed by that ancestral population to the present (Figure 2B,C; Figure S4; see Appendix A.3.2 for an explanation). While ancient individuals were likely not drawn from populations directly ancestral to present-day populations, we should observe the same general patterns with genetic differentiation between the ancient and present-day population (see also Carlson *et al*., 2021, for approximations of this decline).

For the remainder of this article we consider A and B to be contemporary populations; we assume for simplicity that they split from a common ancestral population without subsequent gene flow. In Figure 3A, we use simulations and analytical predictions to show how the prediction accuracy in population B decreases with increasing time since its common ancestor with population A. The span of neutral genetic differentiation was chosen to reflect a scale along which various human populations could fall. We see slightly weaker reductions in prediction accuracy with time when the variance of the mutant effect size distribution is quartered and stronger reductions when the mutant effect size distribution has a heavy tail (Figure 3B; Figure S5). All variance-contributing loci were ascertained in population A, so the only reason the full genic variance in population B was not captured is because private polymorphisms contribute to the phenotypic variance in each population. At the time of divergence between the pair of populations, they entirely share their genetic basis of trait variation. Then as stabilizing selection and drift remove polymorphisms (at equilibrium), new mutations replenish them at different sites in each population, leading to the same total variance but different genetic bases of that variance (assuming a very large mutational target). Without gene flow between the pair of populations, the polymorphisms that arose since their common ancestor will remain private to each population and will thus not contribute to the variance in the other population. Therefore the polymorphisms in population A that also contribute to the variance in population B must be ancestrally shared polymorphisms, specifically at those loci in which ancestral polymorphisms were not removed by drift or selection in either descendant population. Note that these shared polymorphisms on average contribute less variance in the descendant populations than in their ancestor. Thus we see a loss in the additive genetic variance shared between populations over time due to a reduction in both the number of shared ancestral polymorphisms and the variance they each contribute. In Appendix A.3.2 we describe our analytical predictions for this process, which match well to simulations.

**Figure 3:**
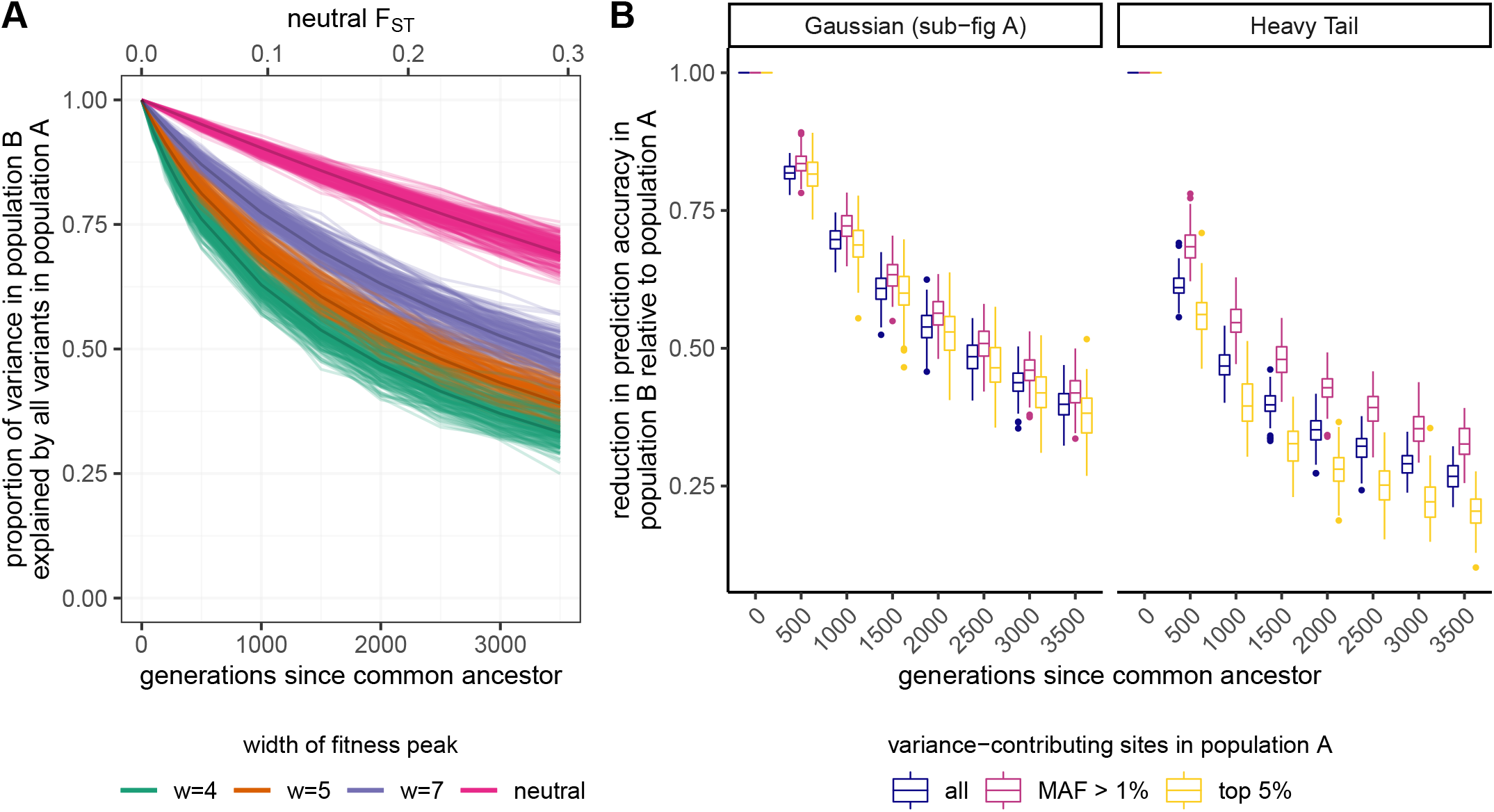
**A)** Reduction of prediction accuracy in population B, when all variance-contributing polymorphisms in population A were ascertained, with increasing time since the common ancestor of populations A and B. Lighter lines show results from simulations and darker lines show results from analytical predictions. **B)** Distribution of the reduction in prediction accuracy for population B relative to population A from simulations with *w* = 5 when different sets of variance-contributing sites in population A were ascertained. Results are shown for two mutant effect size distributions. The Gaussian distribution corresponds to the blue distribution and the Heavy Tail to the yellow distribution in the inset of Figure 1C. The same Gaussian distribution was used to produce results in sub-figure A.

### 4.2 Ascertainment of subset of causal loci in GWAS sample

To more realistically explore the impact of GWAS ascertainment, we explore power- and frequency-based variant discovery. In the main text, we focus on a case in which a GWAS uncovered the top 5% of variance-contributing polymorphisms in population A. Under our choice of effect size distribution and strengths of stabilizing selection, these polymorphisms explain just under 50% of the additive genic variance in population A (similar to that of height in Europeans; Yang *et al*., 2010; Wood *et al*., 2014; Yang *et al*., 2015). We also consider a case in which a GWAS uncovered all causal loci with a minor allele frequency (MAF) that exceeds 1%. As expected, under any ascertainment scheme we observe a substantial drop in the variance explained in population B compared to the case in which all polymorphisms in population A are ascertained (Figure 3B). If this reduction in variance for each population is the same, then the decline in prediction accuracy in population B *relative* to population A over time will be similar to the case when all polymorphisms in population A are ascertained. We find that this is approximately the case for a Gaussian mutant effect size distribution but not for the heavy tail effect size distribution that we simulate (Figure 3C). These differences arise from differences in the strength of selection on and variance contributed by ancestrally shared polymorphisms; see Supplement S1.2 for details. We present results under GxE, pleiotropy, and directional selection in Supplement S1.3.

## 5 Difference in polygenic means among populations

In the previous section we described how stabilizing selection and ascertainment in the GWAS sample can reduce the prediction accuracy of individual genetic values, but what are the consequences for the mean polygenic score of populations? The mean polygenic score of the population is twice the sum of population allele frequencies weighted by effect sizes. If a trait is neutrally evolving, the loci contributing to its variation are just like other neutrally evolving loci, and so differences among populations in their mean polygenic score just reflect a weighted sum of neutral allele frequency differences. Naively, as trait increasing alleles underlying a neutral trait are equally likely to drift up or down, one might think that over many loci we expect only a small mean difference between populations. However, the polygenic score is a sum rather than a mean, and so each locus we add into the score is like an additional step in the random walk that two populations take away from each other (Chakraborty and Nei, 1982). We expect the variance among populations, i.e. the average squared difference between population means and the global mean, to be 2*V*_*A*_*F*_*ST*_ (Wright, 1951; Lande, 1992). We first explore the differentiation of mean polygenic scores under a constant optimum and constant environment, and then relax these assumptions.

### 5.1 Stabilizing selection to a constant optimum

Under a constant selective environment, stabilizing selection keeps population mean phenotypes close to their optimum in the face of genetic drift and mutation, such that the difference in mean phenotypes among populations with the same optimum should be minor relative to neutral expectations (even accounting for the lower genetic variance within populations under stabilizing selection; Figure 4A). This reduction in the divergence of the mean phenotype between populations reflects the fact that if trait-increasing alleles accidentally drift up in frequency, thus pushing the population mean away from its selective optimum, trait-decreasing alleles are subject to directional selection in their favor (and vice versa).

**Figure 4:**
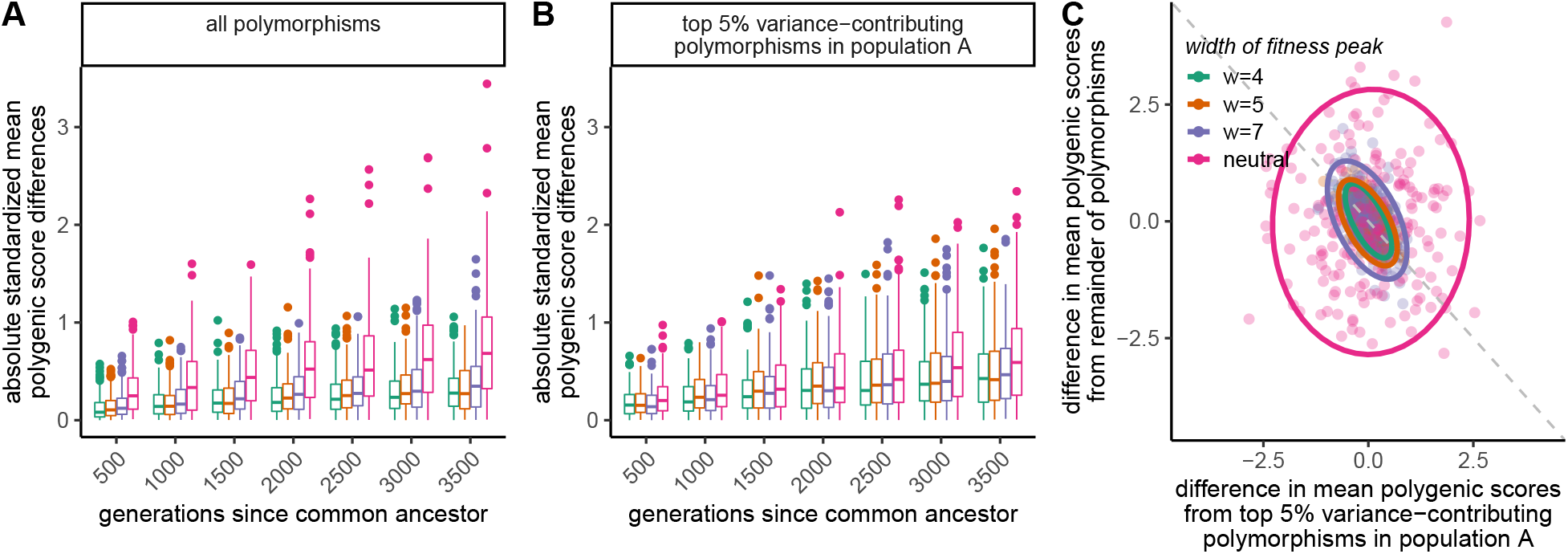
**A)** Absolute standardized mean polygenic score differences between populations A and B, 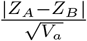, either when all polymorphisms in both populations were ascertained, or when the top 5% of variance-contributing polymorphisms in population A were ascertained. This measure has the same interpretation as *Q*_*X*_ ; it equals 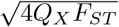. **B)** Partitioning of mean polygenic score differences between the ascertained and non-ascertained set of polymorphisms. Ascertained polymorphisms are from the top 5% of variance-contributing sites in population A. Points represent results for a single simulation. Ellipses denote the 95% confidence interval. The dashed grey line is the line of exactly opposing effects.

To explore population differences in mean polygenic scores, we can use *Q*_*X*_ (Berg and Coop, 2014), *the polygenic score equivalent of Q*_*ST*_*/*2*F*_*ST*_ (Prout and Barker, 1993; *Spitze, 1993). Q*_*ST*_ measures the proportion of total variance in additive genetic values attributable to among-population differences. Under neutrality, for strictly additive traits we expect *Q*_*ST*_ to equal 2*F*_*ST*_ estimated from neutral polymorphisms (Lande, 1992; Whitlock, 1999), and their ratio should be *χ*^2^-distributed with degrees of freedom equal to one fewer than the number of populations (an extension of the Lewontin-Krakauer test; Lewontin and Krakauer, 1973). The polygenic score analog is *Q*_*X*_ (Berg and Coop, 2014), where additive genetic values are substituted by polygenic scores. For our pair of populations, our *Q*_*X*_ statistic is

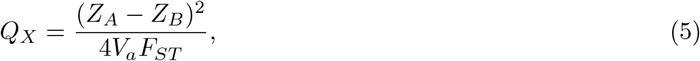

where *Z* is the mean polygenic score in the population denoted in the subscript. When population mean phenotypes are over-dispersed relative to neutral expectations, *Q*_*X*_ will be larger than 1, potentially resulting in a statistically significant p-value under the null distribution. When population means are under-dispersed relative to neutral expectations, as we would expect for traits under stabilizing selection with the same optimum across populations, *Q*_*X*_ will be much smaller than 1, and the p-values under the null will be large. Stabilizing selection to the same optimum tightly constrains the difference in mean additive genetic values between populations, i.e. the difference in mean polygenic scores using all of the variation, to be much lower than the difference for neutral traits even after standardizing for the lower overall levels of variation (Figure 4A). This leads to a distribution of *Q*_*X*_ that is skewed toward lower values than the neutral *χ*^2^-distribution (Figure S12). However, when we ascertain the top 5% of variance-contributing polymorphisms in population A, the level of standardized polygenic score differentiation under stabilizing selection becomes more similar to neutral levels (Figure 4B) with a comparable level of false positive signals of adaptive differentiation as in the neutral case (Figure S12). While this result is more specific to our choice of mutant effect size distribution and strengths of stabilizing selection, we can generally conclude that for cases of stabilizing selection with limited ascertainment, estimates of standardized mean polygenic score differences will approach and perhaps exceed what we observe under neutrality. Note that under the neutral case, the distribution of standardized mean polygenic score differences stays about the same between ascertainment levels, whether ascertaining all polymorphisms from both populations or just the top 5% in population A. This is because under neutrality, all alleles exhibit the same behavior and do not evolve in coordination with one another. Thus stabilizing selection, combined with incomplete and asymmetric ascertainment, causes the inflation of estimated mean standardized population differences above that seen for the underlying genetic values.

Since under stabilizing selection to the same optimum we expect only small differences between populations in their mean genetic value, the disparity between the (true) differences in mean genetic values and (estimated) differences in mean polygenic scores represents the mean difference contributed by non-ascertained polymorphisms. Intuitively this occurs because if population B has a larger value of an ascertained polygenic score than population A, then the ascertained trait-increasing alleles have by chance drifted up in population B (compared to A) and this imbalance will have induced directional selection for the rest of the trait-increasing alleles to decrease their frequency to keep the population close to the optimum (under high polygenicity). In line with our expectations, we find that the mean polygenic score differences calculated from ascertained sites and mean polygenic score differences calculated from non-ascertained sites are close to opposite one another (Figure 4B). The countervailing effect of the non-ascertained loci is noisier when stabilizing selection is weaker, because with weaker selection population means can drift further from their optimum. These results confirm that mean polygenic scores calculated from all polymorphisms (i.e. additive genetic values) should closely match between populations experiencing stronger and similar selection pressures and highlight how incomplete ascertainment can by chance generate misleading differences between them.

### 5.2 Adapting to a changing optimum or environment

The combination of changes in the stabilizing selection regime and incomplete ascertainment of causal polymorphisms can also generate misleading differences in mean polygenic scores and signals of differential selection among populations. When the optimal phenotype shifts from its ancestral value equally in both descendant populations, this directional selection generates responses in the true mean genetic values in each population that track each other extremely closely. However, because our polygenic score based on ascertained SNPs explains less of the variance in population B than population A, we capture less of its response to selection and thus artificially generate a difference between populations in their mean polygenic scores (Figure 5A). This shift in the mean polygenic score between populations, as well as a decrease in the total variance explained, can push the distribution of *Q*_*X*_ towards larger values than the neutral case (Figure 5B). The chances of getting a p-value below a significance threshold of 0.05 are highest when stabilizing selection is strongest (*w* = 4; up to 30% chance depending on time since the optimum shift) but tend to be greater than neutrality for the strengths of selection we investigated (Figure S6). These signals of polygenic selection are consistent with the idea that *Q*_*X*_ informs us about directional selection on polygenic scores, as selection has driven an increase in the polygenic score of A relative to B. However, they also highlight the very incomplete picture that we obtain about selection and genetic differences in the phenotype.

**Figure 5:**
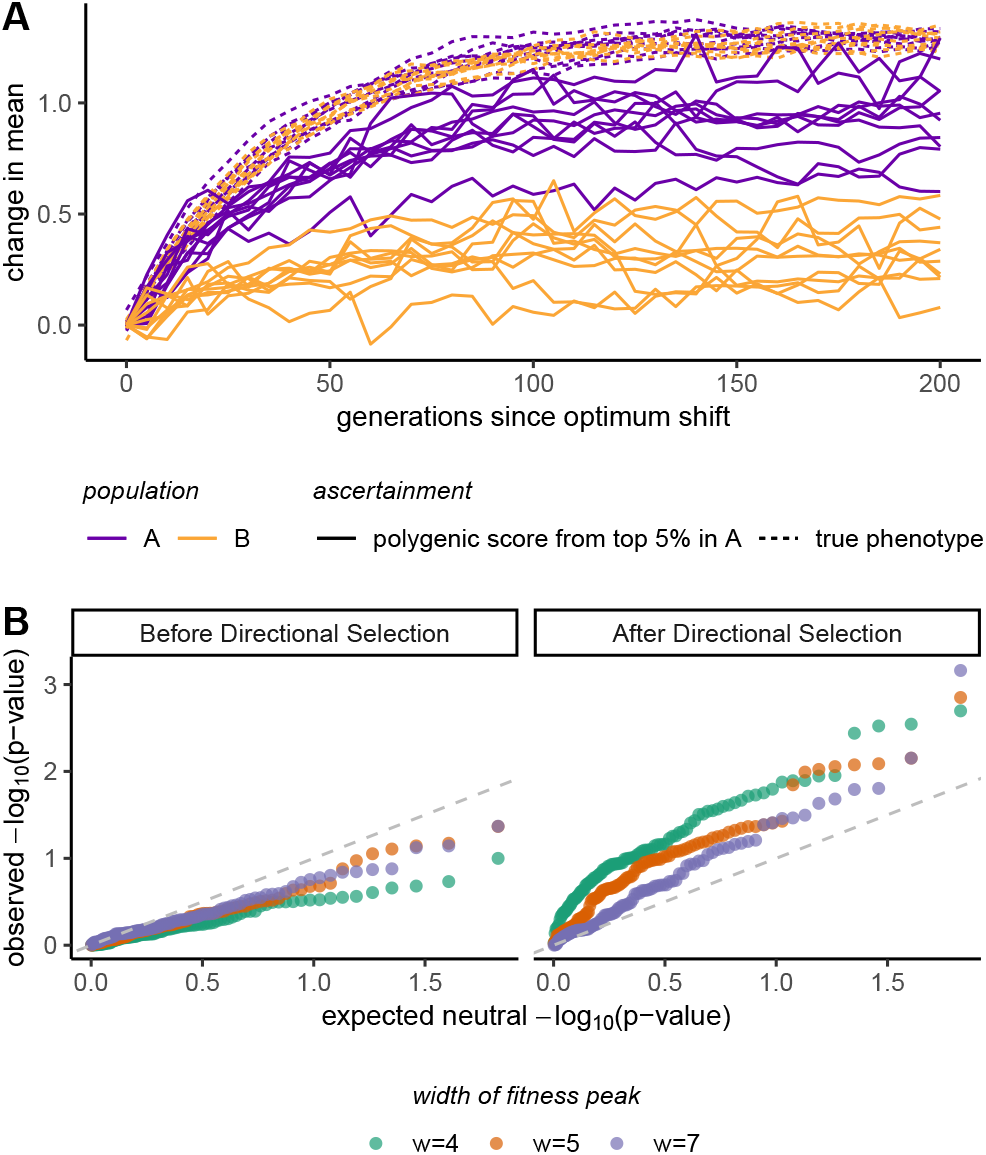
**A)** Change in mean phenotype or polygenic score over time since the optimum shifted 2 standard deviations of the phenotype distribution. True phenotypes are different from additive genetic values in that they account for the contribution of substitutions. **B)** Quantile-quantile plot of observed p-values against expected p-values under neutrality (uniform distribution). The dashed line shows equality; points that lie above this line indicate that the observed distribution has a higher density of low p-values than the neutral distribution, and points that lie below it indicate the opposite.

## 6 Discussion

Our work builds on population genetics theory of stabilizing selection to investigate why polygenic scores perform poorly in unrepresented populations and how they can be misleading about average genetic differences among populations. Specifically, we provide a theoretical foundation on which to understand how stabilizing selection on complex traits impacts the population differentiation of polygenic scores without implicating a difference in environments or trait optima. Negative selection has been invoked as an explanation for low portability consistent with the lack of common, large effect alleles discovered by GWAS (Wang *et al*., 2020; Durvasula and Lohmueller, 2021). Stabilizing selection leads to similar observations, and so our work adds to these recent investigations. Our focus explicitly on a model of quantitative trait evolution allowed us to better inform the relationship between an allele’s phenotypic and fitness effects. By using this approach to model allele frequency dynamics, we can learn more about the relative contribution of population-ascertainment bias on portability, interpretations of genetic differentiation of GWAS loci, and population differentiation of polygenic scores.

### 6.1 Dynamics contributing to low portability

While stabilizing selection with the same optimum among populations reduces phenotypic differentiation relative to neutrality, somewhat counter-intuitively it increases genetic differentiation at trait-influencing loci compared to neutral polymorphisms. Genetic differentiation increases because stabilizing selection drives the turnover of polymorphisms that contribute to trait variation within populations; thus polymorphisms that are ancestrally shared between populations are eventually lost and replaced by private mutations. The consequences of this differentiation at large effect QTLs and few loci have been explored by various authors (Latta, 1998; Le Corre and Kremer, 2003; Kremer and Le Corre, 2012), and here we investigated its implications in the age of GWAS. Over time, the sets of polymorphisms that contribute substantial variance in each population diverge, such that stabilizing selection reduces the additive genic variance in one population that can be explained by polymorphisms in another. Thus polygenic scores constructed from polymorphisms ascertained in a GWAS will have reduced prediction accuracy in unrepresented populations compared to represented ones. The prediction accuracy for unrepresented populations decreases with increasing strengths of stabilizing selection and time since the common ancestor with represented populations. Occasional fluctuations in the fitness optima would negligibly influence our portability results, because such directional selection would cause very minor shifts in frequency over the short time scales we consider (Hayward and Sella, 2021).

The distribution of mutant effect sizes is critical to understanding the effects of stabilizing selection on the turnover of genetic variation among populations. With increasing weight towards larger mutant effects, stabilizing selection will cause a faster reduction in the shared additive genetic variance, and thus polygenic score portability. In addition, the decline in prediction accuracy for unrepresented populations relative to represented ones will depend on frequency- and power-based ascertainment schemes; since polymorphisms shared between populations will more likely be of small effect and common, the discovery of polymorphisms based on minor allele frequency will lead to a weaker decline than discovery based on variance contributed. Conversely, if the trait was truly infinitesimal, stabilizing selection would not contribute to the loss of heterozygosity, such that the sharing of genetic variance would be well predicted by genetic drift, or neutral *F*_*ST*_. While the loci mapped by GWAS are often of small effect, and the traits highly polygenic, we know that their effects are not infinitesimal, as loci must make a reasonable contribution to the variance to be discovered by a GWAS (see Simons *et al*., 2018, for a detailed population genetic model).

While we do not consider migration between represented and unrepresented populations here, the general decline in portability with increasing genetic differentiation between groups should hold, though the exact prediction will differ. Pleiotropy and GxE complicate the predictions of stabilizing selection for portability, which we discuss in Supplement S1.3. Differences in the environment among populations can lead to changes in the effect sizes of alleles (GxE), which reduces the portability because (i) effect sizes will be only partially correlated between populations and (ii) stabilizing selection will purge more of the ancestral variation shared between populations since GxE increases the overall trait variance when the interacting environment changes. Pleiotropy, in our implementation, leads to the opposite effect. Holding the average strength of selection on all alleles constant, when an allele can independently affect multiple traits under stabilizing selection, the correlation between its effect size on the trait of interest and its selection coefficient weakens. Thus stabilizing selection purges less of the shared variation and causes a weaker reduction in portability.

### 6.2 Considerations for applications of GWAS results across groups

Like much of the recent work on portability, our work emphasizes the reduction in prediction accuracy for populations not represented in GWAS (Márquez-Luna, 2017; Martin *et al*., 2017a; Duncan *et al*., 2019; Lam *et al*., 2019; Weissbrod *et al*., 2021). A natural conclusion is that polygenic predictions that work well across populations will require GWAS across a range of diverse ancestries, in line with other calls to reduce Eurocentrism in GWAS (Li and Keating, 2014; Popejoy and Fullerton, 2016; Martin *et al*., 2019). In addition to changes in allele frequency and linkage disequilibrium, the relationship of polygenic scores to phenotypes will vary across populations due to variation in genetic effects, assortative mating, and differences in GxG and GxE. Indeed the prediction accuracy of polygenic scores for some traits was recently shown to be quite variable across different groups within an ancestry, suggesting that GxE and environmental variation are quite prevalent for some traits (Mostafavi *et al*., 2020). In addition, associated variants on the same haplotype were found to have differing effects in white and Black Americans, suggesting genetic or environmental interactions modify additive effect sizes across groups (Patel *et al*., 2021). Thus, we caution that a better understanding of the portability of polygenic scores across populations will also need a stronger understanding of the causes of variation in prediction accuracy within populations.

With the rise of ancient DNA sequencing, GWAS from contemporary populations have also been used to construct polygenic scores for ancient individuals (reviewed in Irving-Pease *et al*., 2021). These scores have been used to provide a window into the phenotypic diversity of past populations (Mathieson *et al*., 2015; Berens *et al*., 2017; Martiniano *et al*., 2017) and to disentangle genetic and environmental contributors to temporal phenotypic variation (e.g. at the Neolithic transition; Cox *et al*., 2019, 2021; Marciniak *et al*., 2021). Such studies are most convincing when there are relevant phenotypic measurements on at least some ancient individuals and polygenic prediction accuracies can be judged. However, investigators will often not have this luxury, leaving unclear the insight these approaches can provide. Some studies using ancient DNA have identified reduced rates of disease alleles in the past compared to present-day populations. While we have focused here on quantitative traits, rather than disease traits, we caution that purifying selection against risk alleles will lead modern day populations to systematically underrepresent the diversity of disease alleles in the past (Berens *et al*., 2017; Aris-Brosou, 2019; Esteller-Cucala *et al*., 2020; Simonti and Lachance, 2021).

### 6.3 Misinterpretations of group differences based on polygenic scores

When using polygenic scores constructed from the ascertained set of polymorphisms, we increase the possibility of generating misleading signals of differentiation between populations. Stabilizing selection to a constant optimum alone does not generate more false signals of directional selection than what we expect under the neutral evolution of complex traits. However, the standardized difference between populations will be systematically over-estimated with ascertained polygenic scores, have standardized for the lower proportion of the genetic variance is explained. We see this result because with stabilizing selection, the unascertained portion of the variance tends to act exactly counter to the trend seen in the ascertained portion. Misleading signals of adaptive polygenic differentiation can also be generated when the stabilizing selection regime shifts in the same way in each population (such that there is still minor phenotypic differentiation between them) as the ascertained polymorphisms only capture a shift in the ascertainment population. This issue arises because stabilizing selection lowers the proportion of the additive genic variance explained in the non-GWAS population and so we capture a lower proportion of that population’s response to directional selection. Thus this issue of missing the parallel adaptive response across ancestries can be expected in many situations with imperfect portability. Such signals of polygenic adaptation can be useful as *Q*_*X*_ is correctly detecting that directional selection has acted on the genetic variation along the branch leading to the GWAS population, but the signal is very open to the misinterpretation that the non-GWAS population has not also responded to the same selection pressures.

Many traits have likely experienced a mixture of stabilizing selection and bursts of directional selection across human history. Even if the populations share the same phenotypic optimum, if an environmental change systematically shifts one population away from this optimum, there would be directional polygenic adaptation to move that population back towards the optimum, resulting in a difference in polygenic scores but no difference in the mean phenotypes between populations (Harpak and Przeworski, 2021). Therefore, under current ascertainment schemes and pervasive stabilizing selection, the difference in mean polygenic scores among populations provides very unreliable information about the potential role of selection in generating phenotypic differences among populations.

A polygenic score is a prediction of how an individual’s phenotype is expected to deviate away from the sample mean given their genotype at some (large) number of polymorphisms. Sometimes they are quoted as absolute values, but that is always based on an empirical phenotypic mean. The mean polygenic score in a population cannot inform us of the mean phenotype in a population, even if it was constructed from all polymorphisms and averaged across all individuals. This is because these scores are based on polymorphic genotypes alone whereas the absolute phenotype of an individual represents the end product of the entire genome and environment played out through a vast number of developmental processes. Yet it is easy to fall into the trap of believing that a difference in polygenic scores between groups is a strong statement about the difference in the mean phenotype of those groups who also differ in a myriad of environmental and cultural factors (a related set of issues are present in epidemology; Rose, 2001). Thus while the field of human genetics is increasing its power to predict phenotypic variance among individuals within groups, it remains a poor guide to the causes of phenotypic variance among groups with greater environmental and genetic differentiation (Lewontin, 1974; Feldman and Lewontin, 1975).

## 7 Data Availability

Code used to generate simulations and process output can be found at https://github.com/SivanYair/SLiMsims_StabilizingSelection.

## 8 Acknowledgements

We thank Doc Edge, Molly Przeworski, Guy Sella, Michael Turelli, Vince Buffalo, Sohini Ramachandran, Yuval Simons and two anonymous reviewers for their valuable comments on earlier drafts of our manuscript. We also thank the members of the Coop lab for helpful discussions. Funding was provided by the National Institute of General Medical Sciences of the National Institutes of Health (NIH R01 GM108779 and R35 GM136290, awarded to GC).

## A Appendix

### A.1 Simulation Details

In SLiM version 3.6 (Haller and Messer, 2019), we simulated stabilizing selection on additive traits and divergence between a pair of populations with the same optimal phenotype (set at zero). Each simulation consisted of a constant population size of 10,000 diploid individuals and genomes comprising 4000 1bp quantitative trait loci and 1000 1bp neutral loci, with free recombination between loci. Mutations arose at rate 2 × 10^*−*6^ per base pair and for quantitative trait loci their effect size was randomly assigned from a normal distribution or a mixture of normal distributions. We used a Gaussian fitness function setting the fitness of an individual to

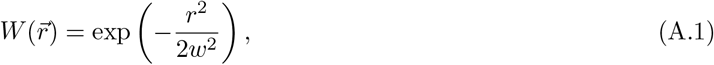

where 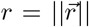 is the phenotypic distance from the optimum of an individual with a vector of phenotypes 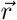 and *w* quantifies the width of the fitness peak. The form of Gaussian selection is consistent with any form of quadratic selection if the population mean is near the optimum (see section A.1.2). We simulated quantitative traits under neutrality (*w* = ∞) and under varying widths of the fitness peak, *w*, in which larger values of *w* indicate weaker stabilizing selection. We chose strengths of stabilizing selection and an effect size distribution that correspond to observations in human populations (see section A.1.1 and A.1.2 for details). We calculated an individual’s fitness each generation based on equation A.1, where their phenotype was the sum of the effects of alleles they carried at both variable and fixed sites.

We burned in each simulation for 60,000 (6*N*) generations before splitting the population into two for 3500 generations. We recorded mutation effect sizes and frequencies in each population every 100 generations for the first 500 generations, and then every 500 generations. To reduce computation times, groups of 10 simulation replicates shared the first 40,000 generations of their burn-in. The remaining 20,000 generations provided more than enough time for a complete turnover in the genetic basis of trait variation, such that simulations within a group were effectively independent. We simulated 100 or 200 replicates of each parameter combination.

#### A.1.1 Choice of parameter values for strength of selection on the trait

We simulated under four different strengths of stabilizing selection (*w*), chosen to range from no selection (neutral trait) to the highest strength of stabilizing selection estimated from the UK BioBank (Sanjak *et al*., 2018). These estimates were determined from regressions of relative fitness (based on estimates of lifetime reproductive success) on squared standard normal phenotypes, *r*^2^, where half of the coefficient of the quadratic term is the quadratic selection gradient, *γ*, in which negative values imply stabilizing selection, and the intercept is the mean fitness in the population, 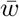. The authors report estimates of *γ* from their regressions, and so to get strengths of stabilizing selection (*w*) under our slightly different fitness model, we make the following approximations to fitness. At equilibrium, the distribution of phenotypes are closely centered around the optimal phenotype relative to the width of the fitness gradient, and therefore we assume that mean fitness 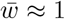. Additionally assuming that the mean phenotype in the population is the optimal phenotype, we can then approximate that the quadratic fitness function that is equivalent to ours when 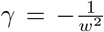, because 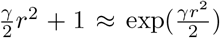. Based on incremental *γ* values of 0, -0.02, -0.04, and -0.06 (the maximum estimated), we use the following values of the strength of stabilizing selection on the trait: *w* = ∞ (neutral trait), *w* = 7, *w* = 5, and *w* = 4.

#### A.1.2 Choice of effect size distribution

Effect sizes of new mutations were drawn from a normal distribution or mixture of normal distributions with mean zero. Provided that the strengths of selection we chose correspond to selection on standard normal phenotypes, we used a standard deviation that would result in a genetic variance close to 1 at equilibrium. However, we note that the size of the mutational target provides a simple inflation factor to the variance and cancels out when calculating the reduction in prediction accuracy; the mutant effect size distribution that we use is more important for determining the distribution of selection coefficients. See equation A.13 for the solution to the equilibrium genic variance. Since the strength of selection helps determine the equilibrium genic variance, by changing the variance of effect sizes of incoming mutations, we used the weakest strength of selection to determine the variance of this effect size distribution (*w* = 7). For a Gaussian distribution of effect sizes, this leads to a solution for the standard deviation (*σ*_*a*_) of 0.1. This distribution generates mutations that are mostly nearly neutral or weakly selected. To explore how different selection regimes lead to different declines in prediction accuracy, we also simulated mutant effect size distributions that involve a higher density of nearly neutral mutations or a higher density of strongly selected mutations. We used a Gaussian distribution with *σ*_*a*_ = 0.05 for the former and a mixture of Gaussians for the latter, which we call the “heavy tail” distribution. The heavy tail distribution generates mutations from a Gaussian distribution with *σ*_*a*_ = 0.05 that has mean 0 with probability 0.65 and mean ± 0.25 with probability 0.175. We chose these specifications for the heavy tail distribution in order to match the median absolute deviation of selection coefficients for the standard Gaussian with *σ*_*a*_ = 0.1.

#### A.1.3 Extension: Directional Selection

We extended our baseline stabilizing selection scenario for directional selection by imposing a positive optimum shift of either one or two standard deviations of the phenotypic distribution. The optimum shifted in the same way 2500 generations after the pair of populations diverged. We started directional selection simulations from the states of the baseline scenario that were recorded at the time of the optimum shift. To calculate the extent of the optimum shift in each simulation, we averaged the phenotypic standard deviation of each population in that simulation. Simulation details and results for other extensions can be found in Supplement S1.3.

### A.2 Relative prediction accuracy of polygenic scores

The prediction accuracy of a polygenic score is the the proportion of the phenotypic variance explained by that polygenic score (the squared correlation *R*^2^). We write out the correlation between an individual’s polygenic score constructed from unlinked polymorphisms discovered by GWAS, *S*_*i*_, and the true additive genetic value, *G*_*i*_, in a particular target population. We assume that a GWAS discovers associations only at causal loci, and that at these loci with significant associations, it estimates their true effects. There are *L*_*s*_ loci with significant associations in the study and a remaining *L*_*r*_ that contribute to trait variation in a particular target population. Therefore an individual’s predicted and true genetic values are

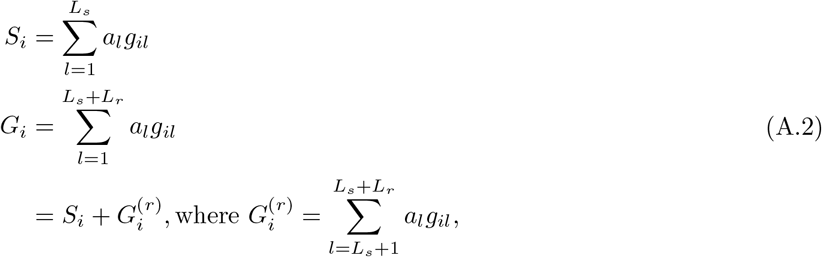

in which their true genetic value is a sum of their predicted genetic value and genetic value contributed by the remainder of sites contributing to trait variation. Because the predicted genetic value is a portion of the true genetic value, the covariance between them is the variance in predicted genetic values,

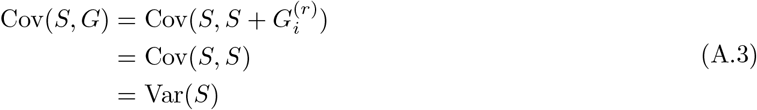

Thus the correlation between the predicted and true genetic values is

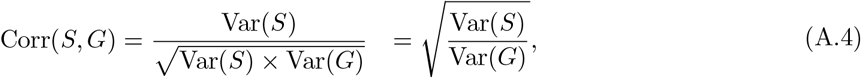

meaning that the correlation between the predicted and true additive genetic values is the square root of the proportion of the trait variance explained by SNPs ascertained by the GWAS, or that the variance in true genetic values explained by polygenic scores (*r*^2^) is the proportion of the variance explained by the polymorphisms used to construct the polygenic scores. This value also represents the reduction in the prediction accuracy of polygenic scores compared to additive genetic values, when predicting true additive phenotypes (with contributions from both genetics and environment). We denote the additive environmental contribution to the phenotype in an individual as *E*_*i*_. The prediction accuracy of an individual’s full additive phenotype (*G*_*i*_ + *E*_*i*_) when using polygenic scores is

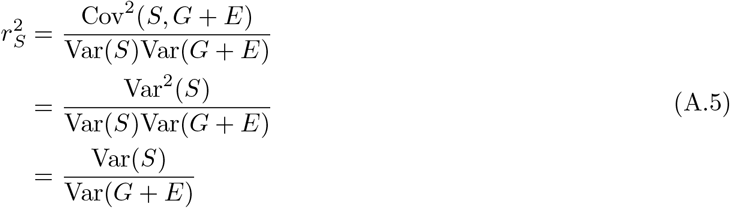

and when using additive genetic values is

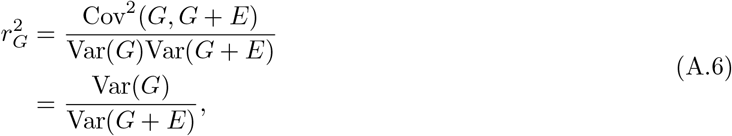

such that the reduction of prediction accuracy when using polygenic scores instead of additive genetic values is

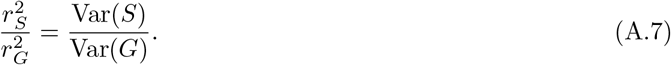

Thus 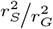 represents the prediction accuracy of polygenic scores for additive genetic values, as well as the reduction in prediction accuracy for full additive phenotypes due to incomplete ascertainment.

#### A.2.1 Modification for gene-by-environment interactions

While we assume the effects of a causal allele are perfectly estimated in the GWAS sample, these effects may differ in populations that experience different environments (GxE). Thus for populations not represented in the GWAS, the correlation between polygenic scores and true additive genetic values will be lower than without GxE. Imagine that we use the set of effect sizes from GWAS population *A* (*a*_*l,A*_) to construct polygenic scores for population *B* where the true effect sizes (*a*_*l,B*_) differ. The covariance of the polygenic scores with true additive genetic values is then

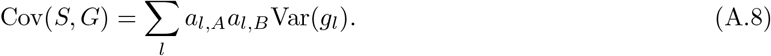

Therefore the correlation between polygenic scores and additive genetic values is

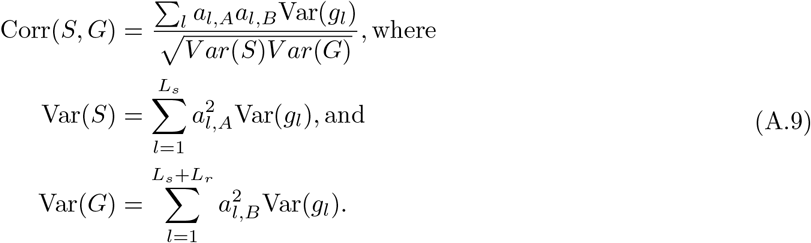

For a neutrally evolving trait, effect sizes and heterozygosities are independent and so

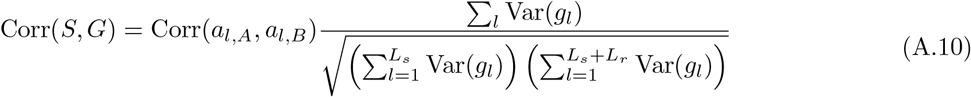

such that the neutral drop in prediction accuracy due to GxE is simply the decrease from 1 of the correlation of effect sizes between A and B.

### A.3 Modeling Details

#### A.3.1 Background

Assuming Gaussian stabilizing selection and that the population mean stays close to its optimum value, selection at an individual locus is well described by a model of underdominance where the per-generation change in an allele’s frequency *x* can be described by its mean and variance,

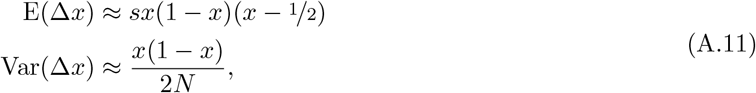

where 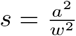 is the the selection coefficient, with *a* being the additive effect of the allele with frequency *x* relative to the alternate allele, and *N* is the diploid population size. This approximation further assumes that the phenotypic variance is much smaller than the width of the stabilizing selection (*σ*^2^ ≪ *w*).

Assuming an infinite sites model, the mean time that a mutation spends in a certain frequency interval before fixation or loss (at a single site) multiplied by its population mutation rate is equivalent to the expected number of alleles segregating at those frequencies at a single time point (see Sawyer and Hartl, 1992). The mutation rate of an allele with effect *a* is 2*NU* Pr(*a*), where *U* is the per-generation mutation rate for the entire mutational target, and Pr(*a*) is the proportion of mutations with effect *a* (with density *f* (*a* | *σ*_*a*_), where *σ*_*a*_ is the standard deviation of the mutant effect size distribution). The mean time *τ* spent in frequency interval (*x, x* + *dx*) for a new mutation with effect *a* can be solved using the Green’s function (see equation A18 of Simons *et al*., 2018), such that

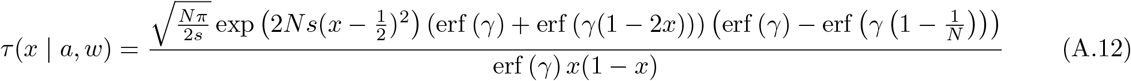

where erf is the error function and 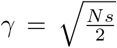. Thus we can solve for the expected total additive genic variance at equilibrium by summing over the heterozygosities contributed by all sites,

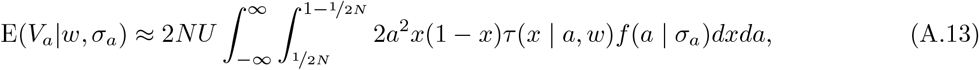

#### A.3.2 Results

In our models we focus on divergence without gene flow between a pair of populations, in which case they only share polymorphisms that arose in their common ancestor and were not lost in either population. To predict the loss of ancestral heterozygosity and probability that the descendant populations share polymorphisms, and thus describe the proportion of additive genic variance in one population explained by polymorphisms identified by a GWAS in the other, we consider the frequency trajectory of ancestrally segregating variants. To fully calculate these quantities we would need the diffusion transition density with underdominant selection to calculate the distribution of the frequency in the current day given the ancestral frequency, but we can build simple approximations of the combined effects of selection and drift. If we assume that the frequency of our allele *x* is close to 0 such that 1 − *x* ≈ 1, then we can approximate its expected per-generation change (eqn (A.11)) as

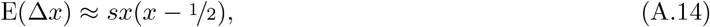

such that its deterministic frequency trajectory follows the logistic function from starting frequency *x*_0_,

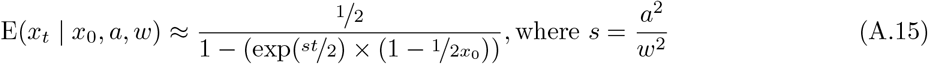

To consider the effects of genetic drift on the frequency trajectory, we assume that *x*_*t*_ is normally distributed around its deterministic frequency trajectory as follows,

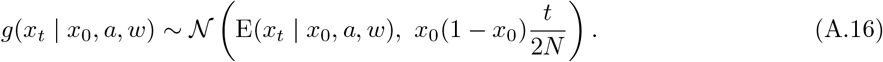

##### Reduction in variance due to drift and selection

We can predict the additive genic variance contributed by all ancestral polymorphisms (anc) to the descendent (desc) after *t* generations by averaging over their effect sizes, starting frequencies, and trajectories,

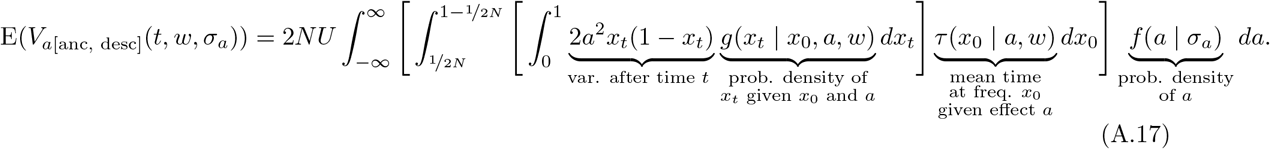

The total reduction in the additive genetic variance contributed by ancestral polymorphisms after time *t* is thus given by 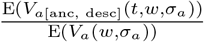.

To obtain the reduction in heterozygosity used in Figure 2A, we use the ratio of expected heterozygosities for the midpoints of the effect size bins evaluated. The equations we use have a similar structure to those where we predict the total variance explained by ancestral polymorphisms at time 0 and *t*, except we do not integrate over all effect sizes and remove *a*^2^ to calculate the heterozygosity instead of variance,

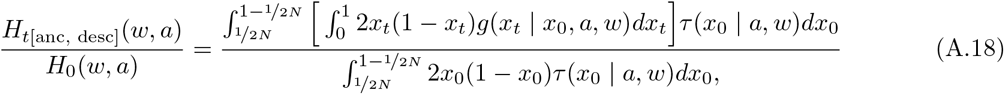

where *H*_*t*_ refers to the heterozygosity at time *t*.

##### Variance in ancestor explained by polymorphisms in descendant

To understand the ancestral variance attributable to present-day polymorphisms, we need to think of the properties of present-day alleles backward in time. For alleles with effect sizes large enough for selection to act strongly, selection (usually) constrains them from reaching appreciable frequencies and selection acts approximately like additive selection against the allele (eqn (A.14)). Conditional on the present-day frequency, the distribution of a deleterious, additive allele’s trajectory backward in time is the same as the process forward in time, such that *g*(*x*_*t*_ |*x*_0_, *a, w*) = *g*(*x*_0_|*x*_*t*_, *a, w*) (Maruyama and Kimura, 1975). Therefore, we can write the expected additive genic variance in the ancestral population explained by variation segregating in the descendant population, in which they have diverged for *t* generations, as

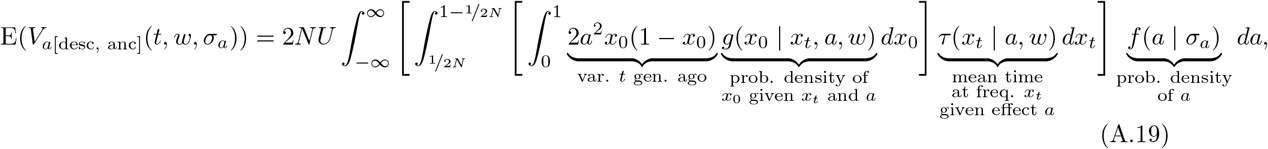

which is identical to the additive genic variance contributed by all ancestral polymorphisms to the descendent (with just a flip in the subscripts, see equation (A.17)). Thus the decay in the additive genic variance segregating in an ancestral population to the present day is the same as loss in prediction accuracy when using present day polymorphisms to predict phenotypes in the ancestral population.

##### Variance in population B explained by polymorphisms in population A

When the population in which we make predictions is contemporary with the population represented in the GWAS, we again consider the variance explained by all polymorphisms that contribute to the variance in the prediction population. In our model of population divergence, these sites that contribute to the variance in the prediction population are sites that were segregating in the common ancestor that remain segregating in both the GWAS and prediction population. We first account for the variance contributed by ancestrally segregating sites in the descendant population in which we make predictions (B), as explained in equation (A.17)), and then for the possibility that each site is segregating, and thus ascertained, in the population from which the GWAS sample was drawn (A),

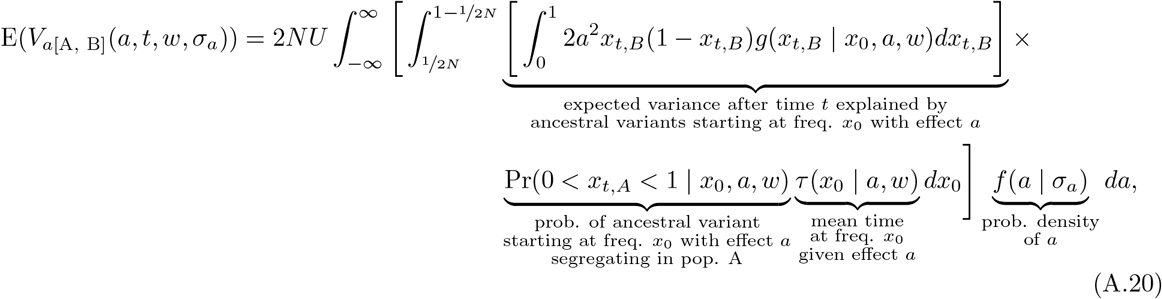

where *x*_0_ is the frequency of an allele in the common ancestor of populations A and B, *x*_*t,A*_ and *x*_*t,B*_ are its frequencies after time *t* in each of those descendant populations respectively, and the probability that the ancestral variant is segregating in population A for ascertainment is given by,

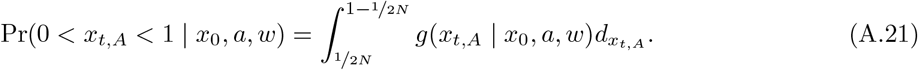

### S1 Supplementary Text and Figures

#### S1.1 Figures referenced in the main text

**Figure S1:**
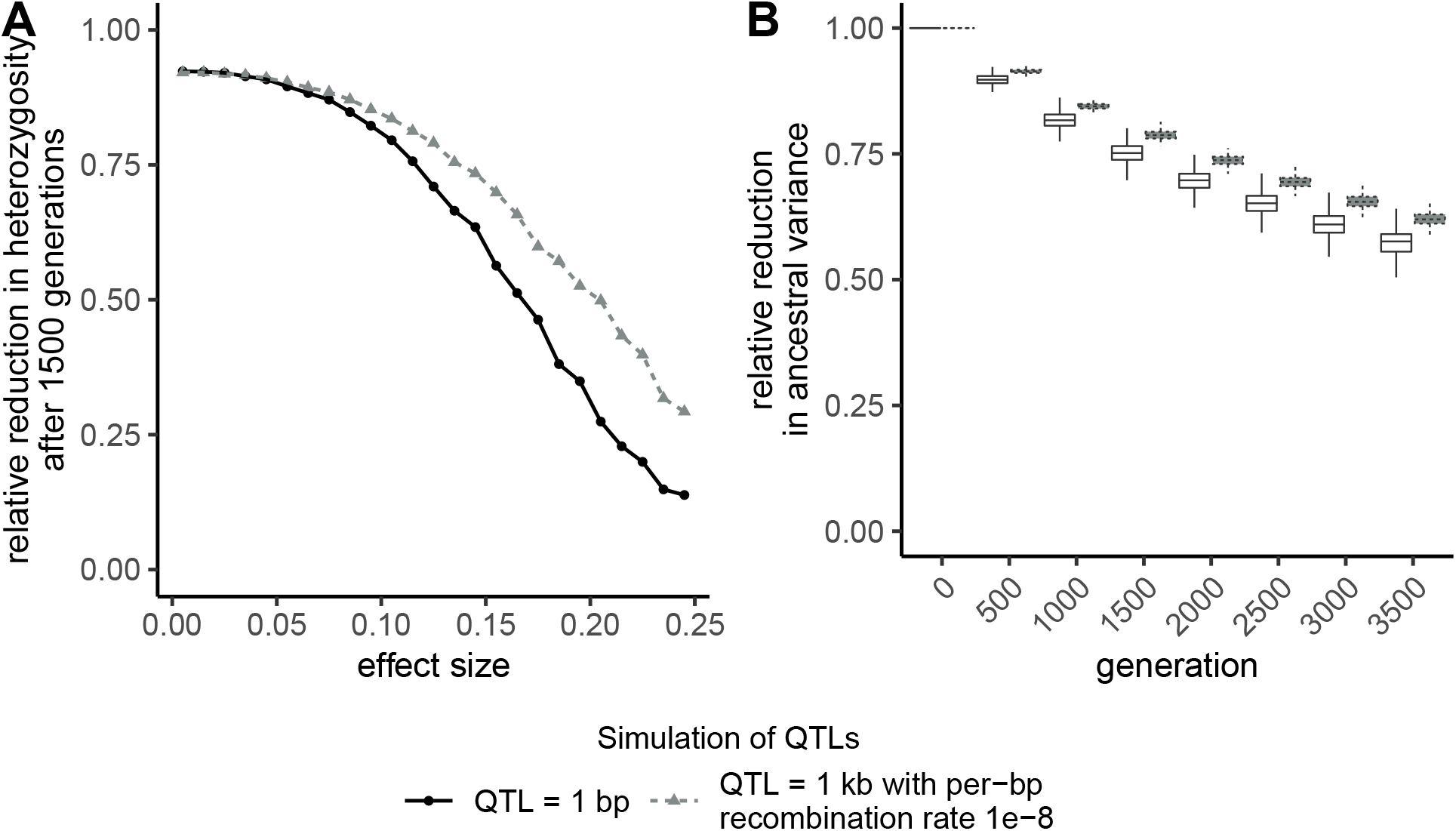
Comparison of **A)** reduction in heterozygosity at loci that contributed to the variance 1500 generations ago within effect size bins of width 0.01 and **B)** total reduction in ancestral variance over time for simulations that differed on the size of QTLs. QTLs were either 1bp (main text results) or 1kb with low within-QTL recombination. Simulations with 1kb QTL consisted of genomes comprising 3000 QTL, with free recombination between QTLs and a per base pair recombination rate of 10^*−*8^ within a QTL. Mutations arose at rate 10^*−*8^ per base pair. All other simulation details match the baseline stabilizing selection scenario introduced in Appendix A.1. The genome-wide mutation rate (*U*) differs for each simulation type, however under the assumptions of our models this should not influence the reduction in heterozygosity or variance.

**Figure S2:**
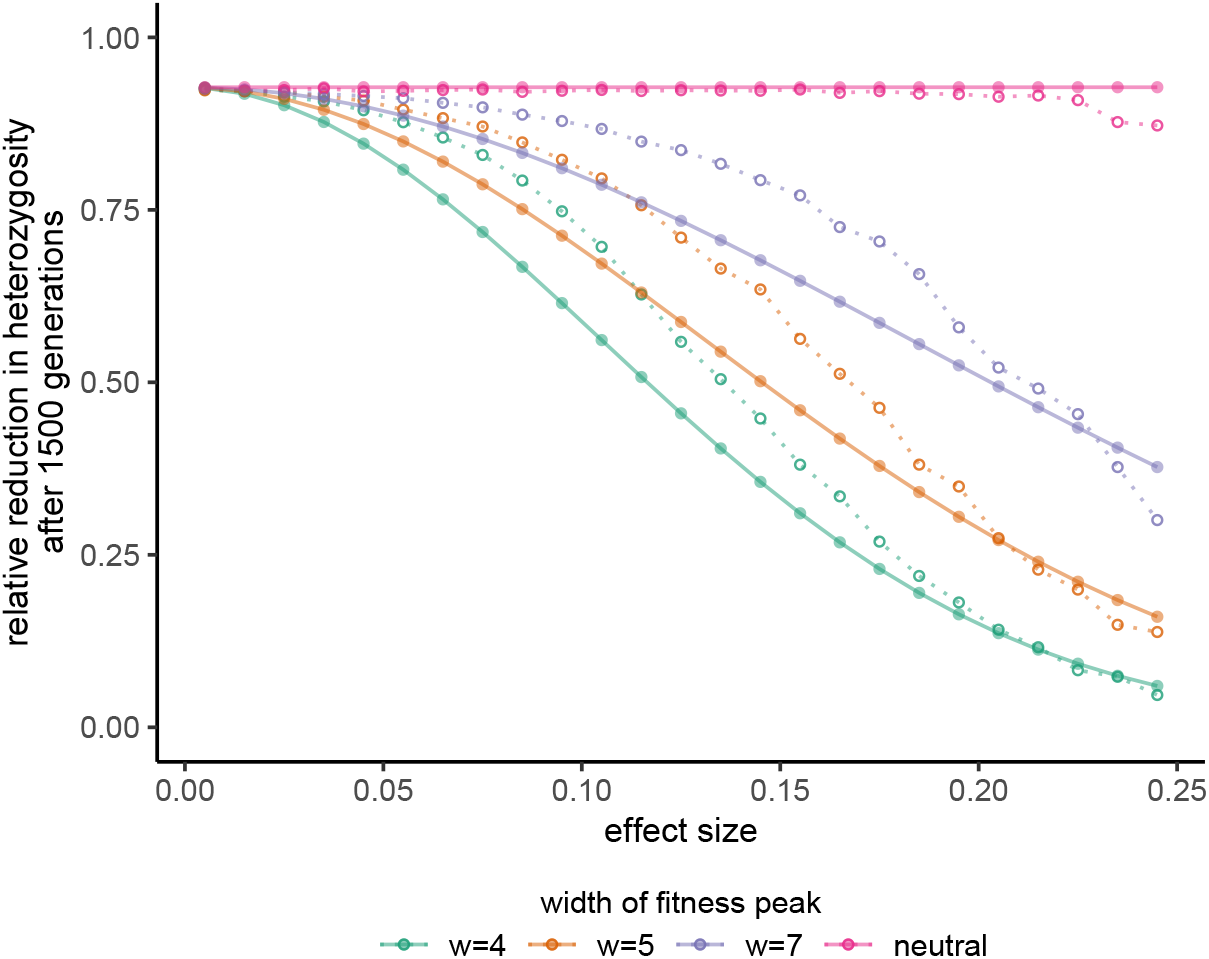
Reduction in heterozygosity at loci that contributed to the variance 1500 generations ago. Each open point, connected by a dashed line, represents the midpoint of the effect size bin of width 0.01 within which we averaged heterozygosity from 200 simulations. Each filled point, connected by a solid line, represents our analytical predictions for that midpoint using the approximation in Equation 3. These predictions over-estimate the reduction in heterozygosity, whereas the predictions from the diffusion approximation provide a much better fit (Figure 2).

**Figure S3:**
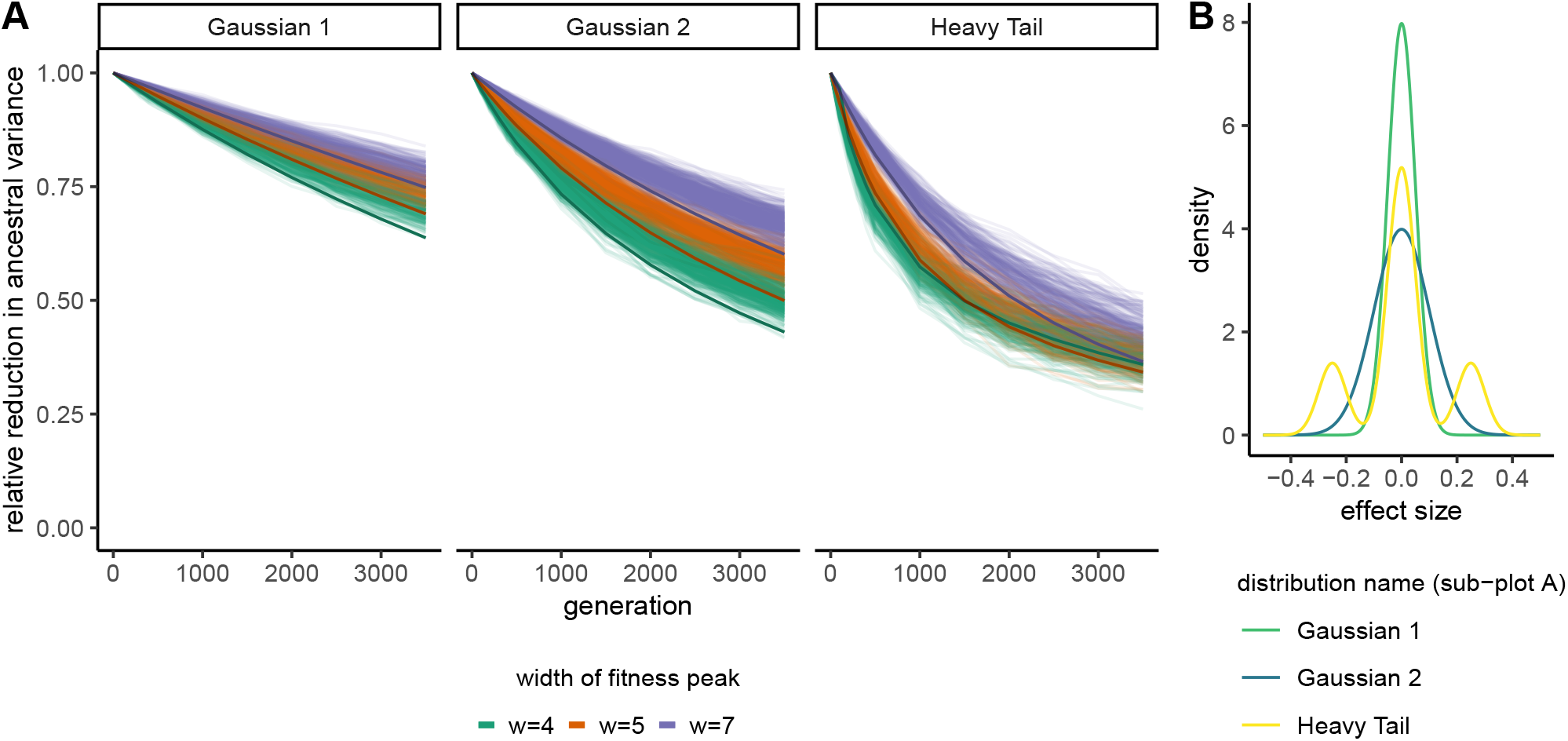
Reduction over time in the total variance contributed by polymorphisms in an ancestral population. Lighter lines show results from simulations and darker lines show analytical predictions based on the diffusion approximation. The results from “Gaussian 2” are the same as presented in Figure 2B.

**Figure S4:**
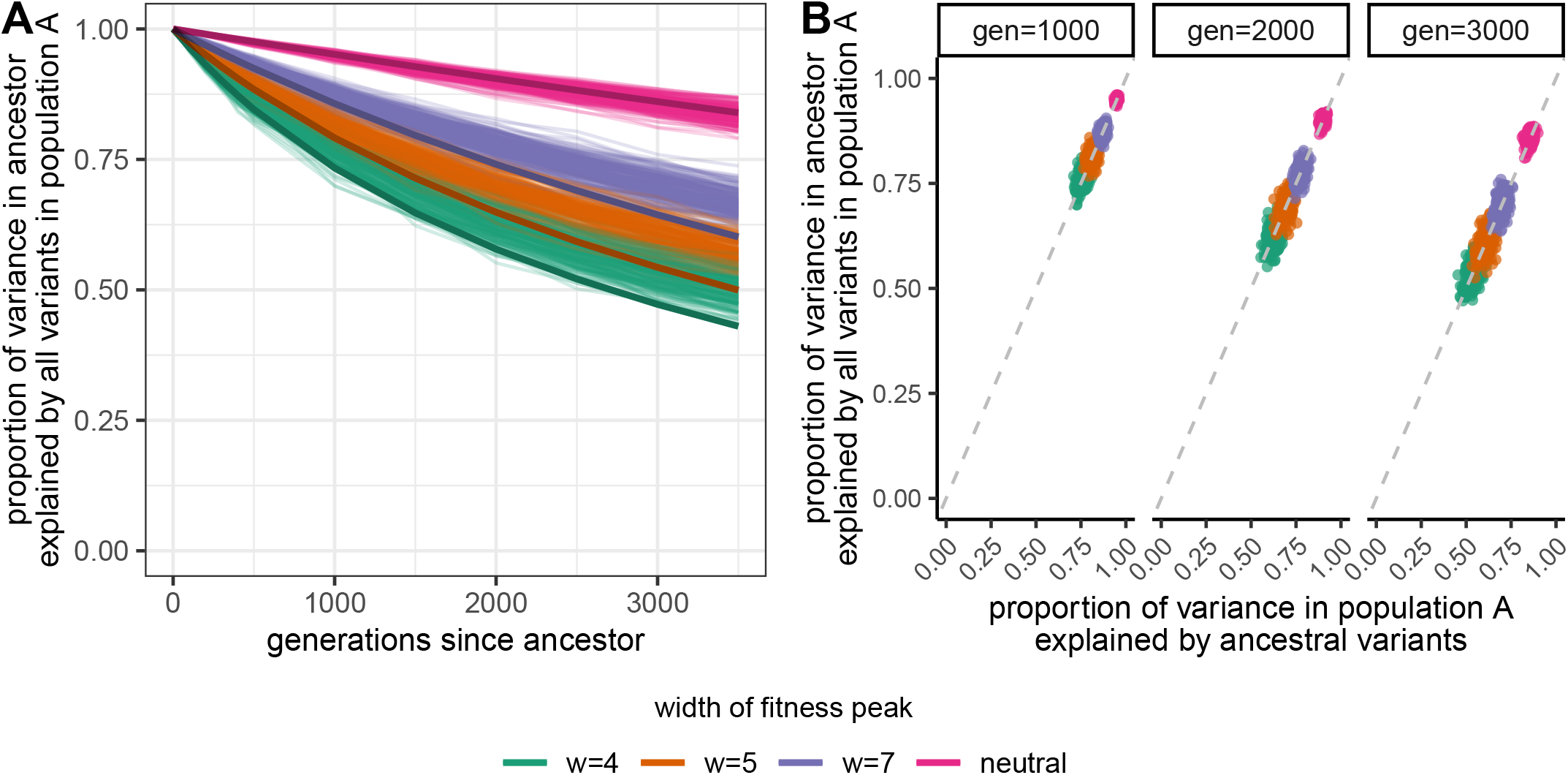
A) Reduction in prediction accuracy in population ancestral to population A with increasing divergence time between them. Lighter lines show results from simulations and darker lines show analytical predictions. B) At particular time depths of divergence, the reduction in prediction accuracy in ancestral population (y-axis) is about the same as the reduction in ancestral variance (x-axis). Points represent results from a single simulation. The dashed line is the line of equality.

**Figure S5:**
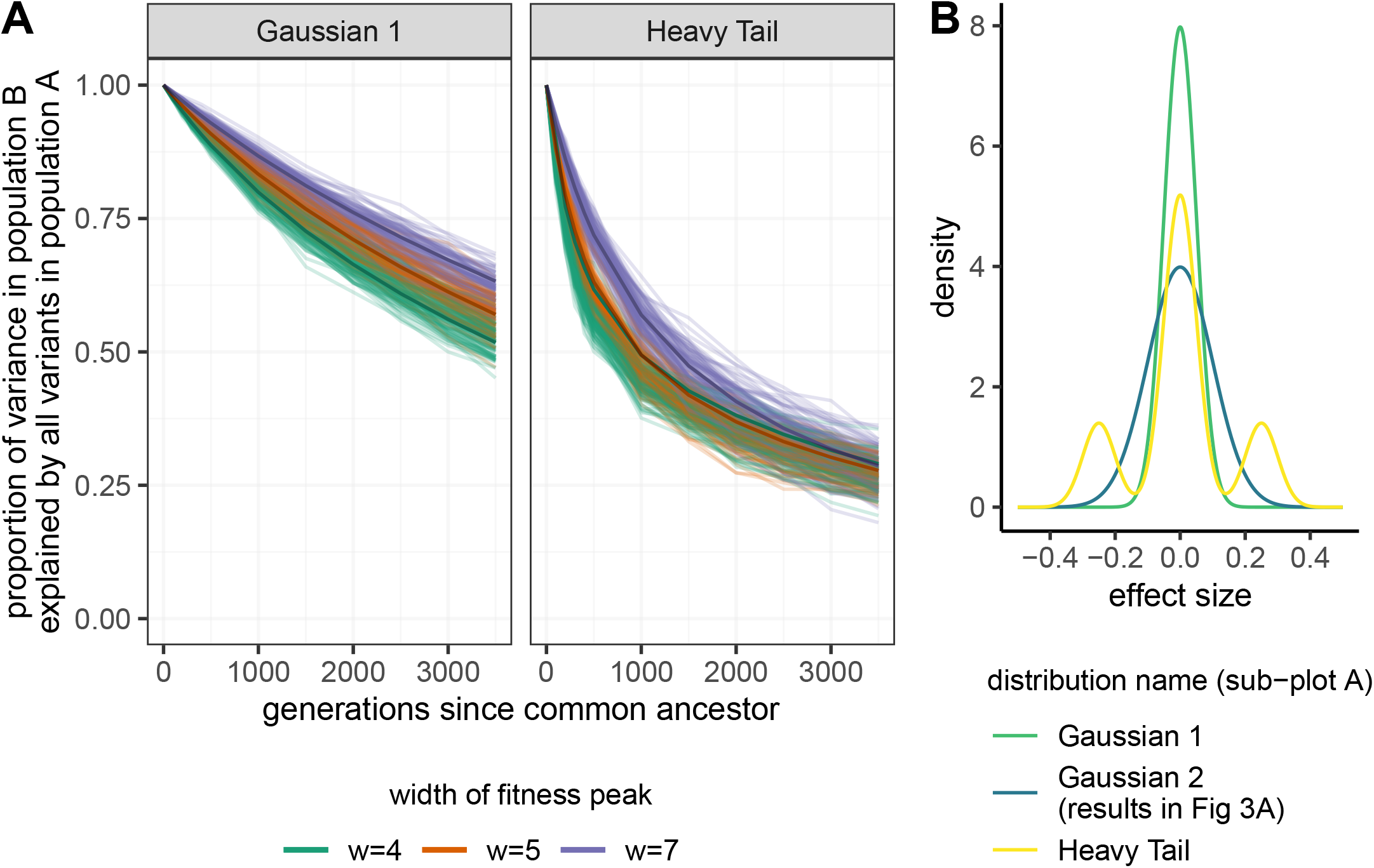
**A)** Reduction in prediction accuracy in population B when ascertaining all polymorphisms in population A, under different mutant effect size distributions than shown in the main text. Light lines show results from simulations, and dark lines show results from analytical predictions. **B)** Mutant effect size distributions correspond to what was used to produce results from sub-figure A. Results from the distribution “Gaussian 2” are shown in the main text Figure 3A.

**Figure S6:**
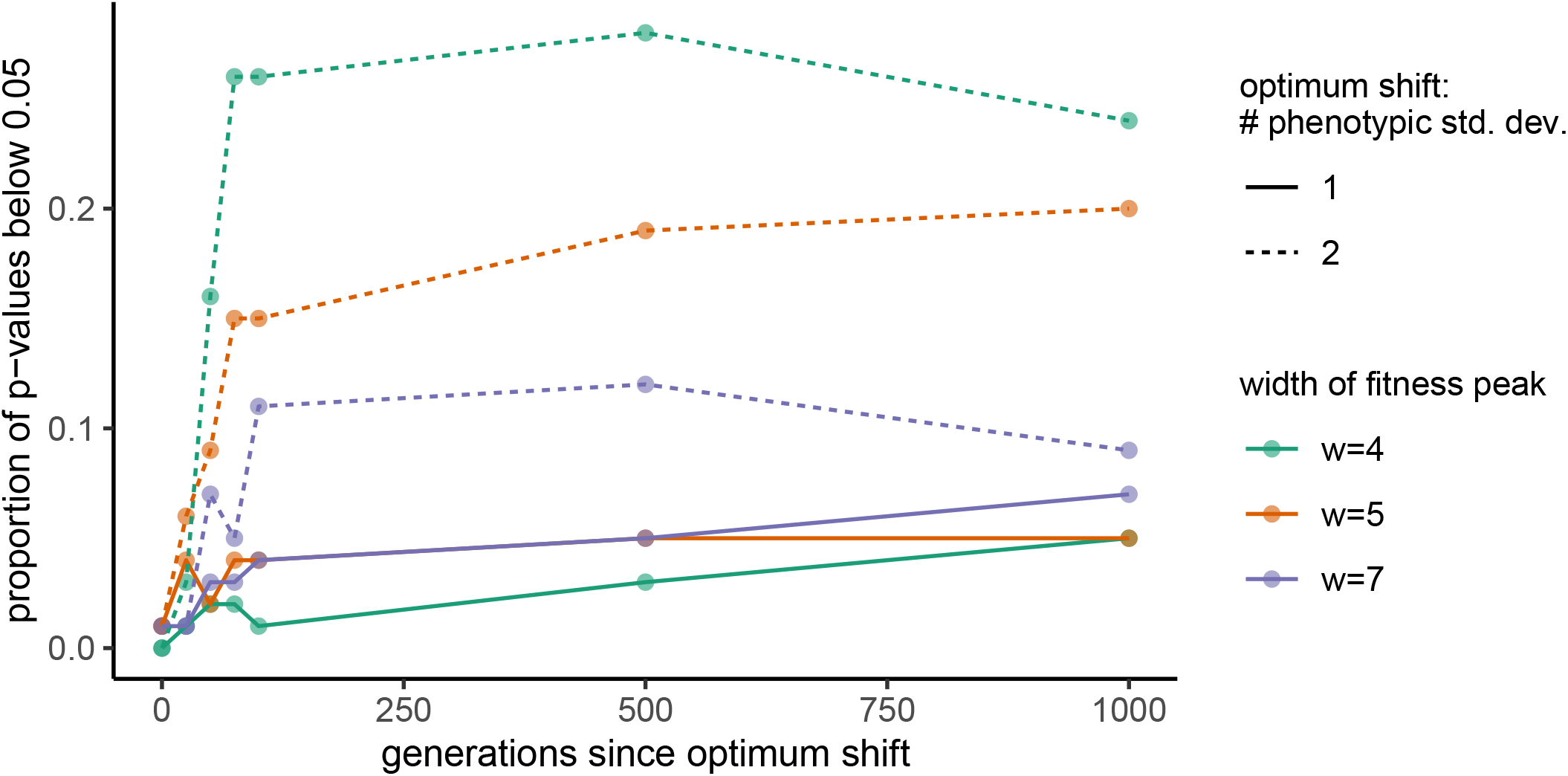
Under the directional selection scenario, proportion of p-values below the significance threshold of 0.05 with time since the shift in the optimum (the shift began at 2500 generations of divergence between populations A and B). Each point is averaged across 100 simulations. Solid lines connect points corresponding to an optimum shift of one standard deviation of the phenotypic distribution, while dashed lines connect points corresponding to an optimum shift of two standard deviations. Following the optimum shift, we recorded results every 25 generations for the first 100 generations, and then every 500 generations.

#### S1.2 Decline in prediction accuracy for population B relative to population A under various ascertainment schemes and mutational effect size distributions

Here we compare the decline in prediction accuracy for population B relative to population A among various ascertainment schemes within population A. Given that any set of polymorphisms discovered within population A is a subset of all polymorphisms in population A, we use the case where all polymorphisms in A were ascertained as the point of comparison for other ascertainment schemes. We simulate cases where populations A and B experienced the same selective environment, such that they have the same expected total additive genic variance at equilibrium. Therefore the prediction accuracy in population B relative to population A is simply the ratio of the variances explained by the GWAS-significant sites in each population. This relative decline will not change when moving from all polymorphisms in A ascertained to a particular subset if the reduction in variance explained for population A and population B is the same. We find that this is approximately the case for a Gaussian mutant effect size distribution containing mostly nearly neutral and weakly selected mutations (Figure 3C; Figure S8). However, the heavy-tailed effect size distribution, which consists of a higher density of strongly selected mutations, leads to different patterns of relative decline for population B when compared to the case when all polymorphisms in A are ascertained. This is because in the heavy-tailed case, most ancestral polymorphisms that are shared between populations are nearly neutral and segregate at intermediate frequencies Figure S7. Thus when moving from ascertaining all polymorphisms in A to just those with MAF*>*1%, population B experiences less of a reduction in variance explained than population A. However, few of those shared polymorphisms explain much of the variance in population A, and thus population B experiences more of a reduction in variance explained when ascertaining the polymorphisms in population A based on variance explained.

**Figure S7:**
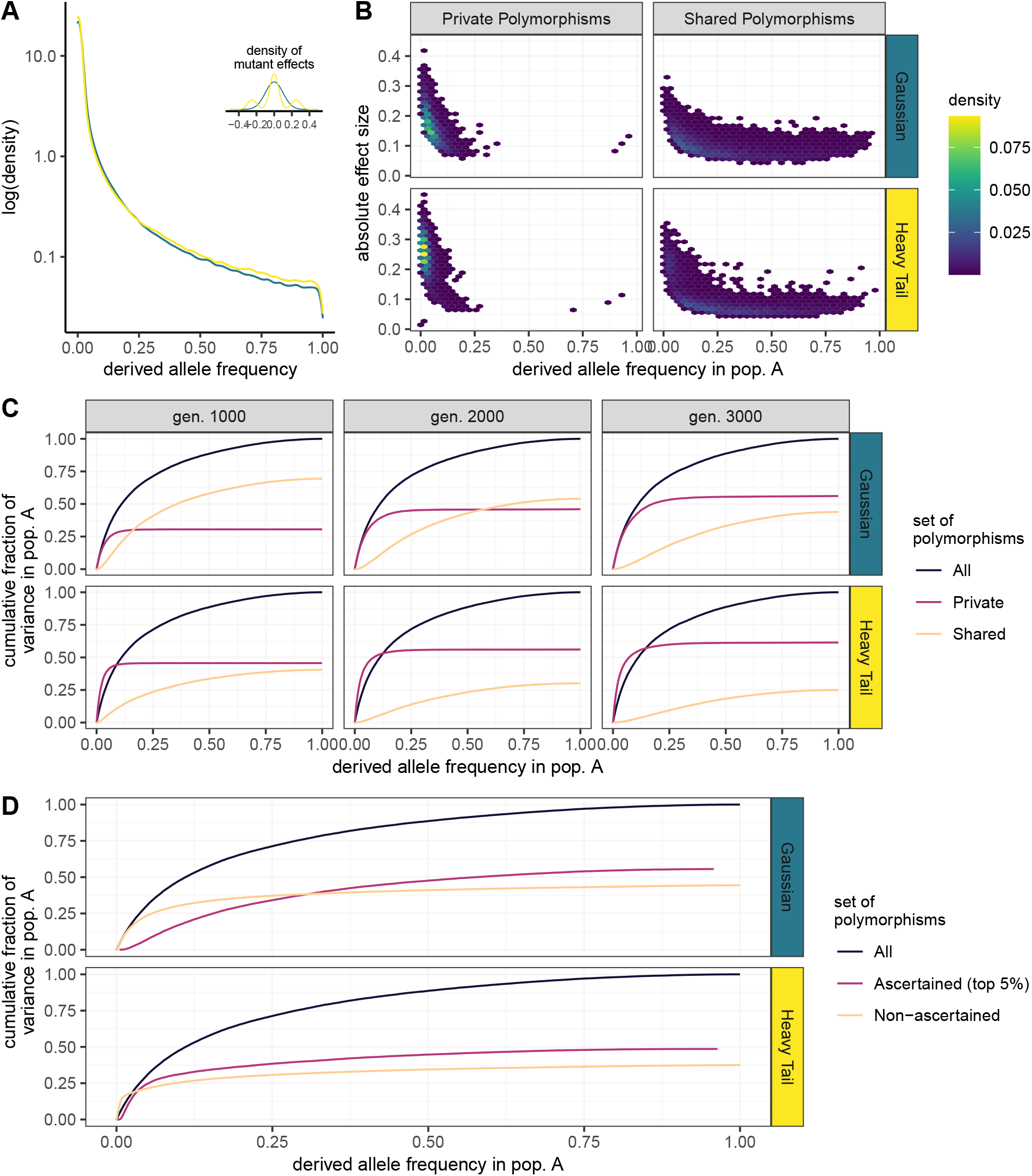
Descriptions of the joint distribution of allele frequencies and effect sizes within a population. **A)** Derived allele frequency spectrum under two mutant effect size distributions shown in the inset, whose colors are used to reference them in the panels of B-D. **B)** Joint density of derived allele frequencies and effect sizes in population A, separated for polymorphisms private to A and shared with B. Note that some private polymorphisms may be polymorphisms in the common ancestor of A and B that persisted in A but were subsequently lost in B. **C)** Cumulative fraction of the total variance in population A with increasing derived allele frequencies in population A for different generations of divergence between population B (columns) and two different mutant effect size distributions (rows). Line colors refer to all polymorphisms in population A, polymorphisms private to population A, and polymorphisms shared between populations A and B. **D)** Cumulative fraction of the total variance in population A with increasing derived allele frequencies in population A for two different mutant effect size distributions (rows). Line colors refer to all polymorphisms in population A, the top 5% of variance-contributing polymorphisms in population A, and the remaining non-ascertained polymorphisms (bottom 95%).

**Figure S8:**
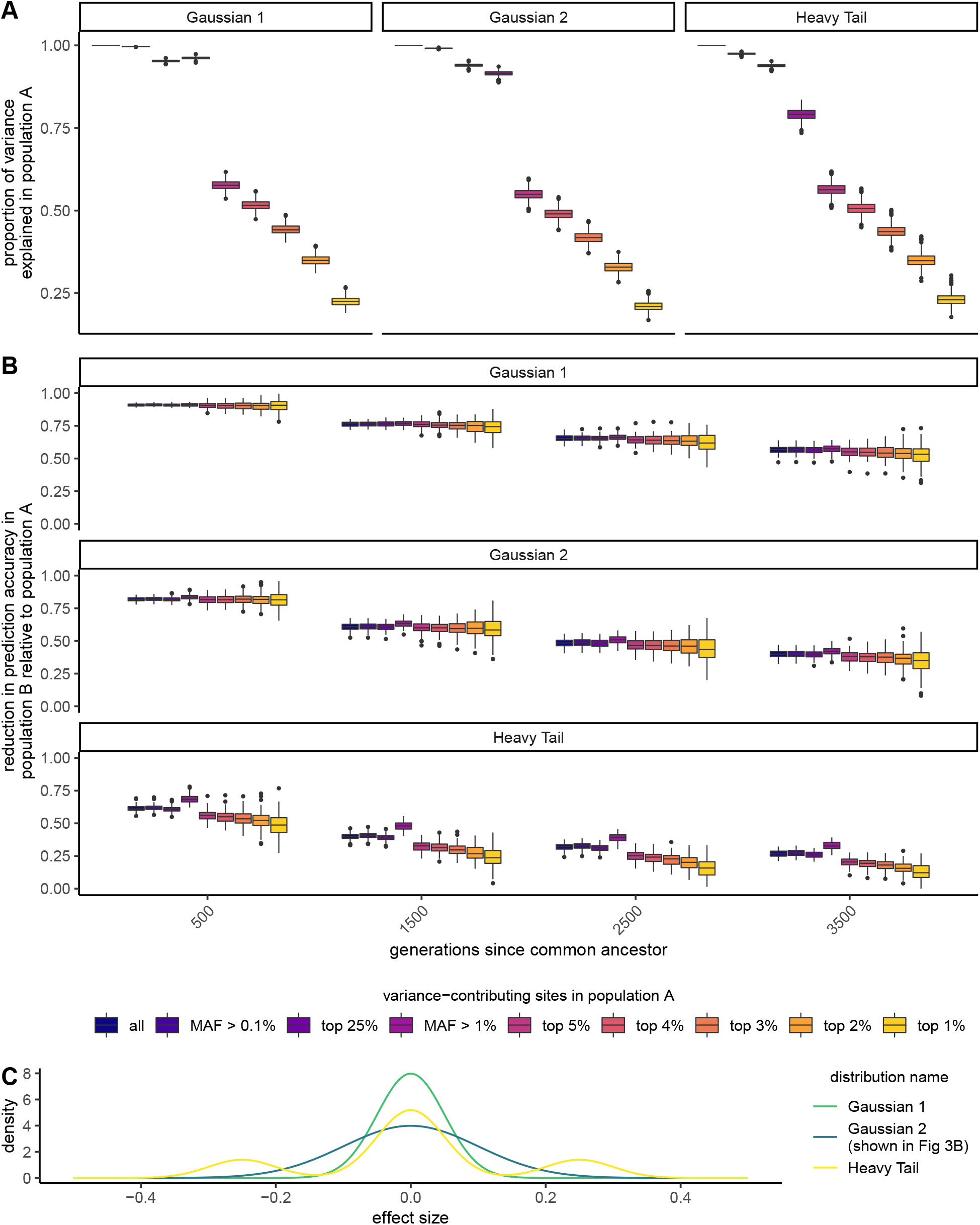
**A)** Proportion of the variance explained in population A when ascertaining different sets of variance-contributing sites in population A. The proportion of the variance explained is the same as the reduction in prediction accuracy for population A when ascertaining a particular set of sites instead of all variance-contributing sites. Facets show results from three different mutant effect size distributions. **B)** Reduction in prediction accuracy for population B *relative* to population A under different ascertainment schemes in population A and mutant effect size distributions. **C)** Mutant effect size distributions that were used to produce results in sub-figures **A** and **B**.

**Figure S9:**
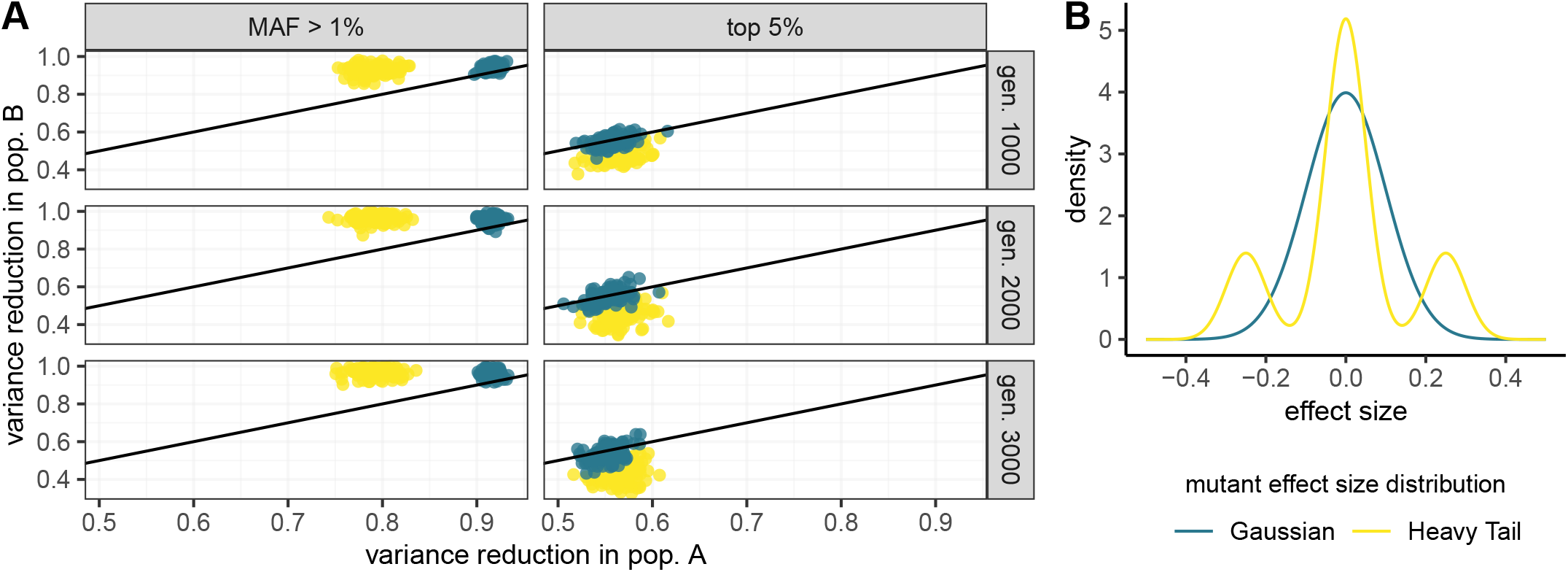
**A)** The relative reduction in the variance explained for populations A and B when switching from ascertaining all variance-contributing polymorphisms in population A to either those polymorphisms with MAF*>*1% or the top 5% of variance-contributing polymorphisms. Points show results from simulations, and their colors correspond to the mutant effect size distributions in **(B)**. The black line is the line of equality. Points that fall along this line indicate the same variance reduction in both populations, and thus a similar decline in prediction accuracy for this ascertainment scheme compared to when all polymorphisms in population A are ascertained. Points above the line indicate that population B experiences less of a reduction in variance explained, and points below it indicate that population B experiences more of a reduction in variance explained, compared to population A.

#### S1.3 Extensions to the baseline stabilizing selection scenario S1.4 Directional Selection

Simulation details for directional selection are provided in Appendix A.1.3. We also discuss in the main text how the same optimum shift among populations can lead to false signals of adaptive differentiation.

##### Accuracy of polygenic score predictions

The observed relative reduction in prediction accuracy for the baseline scenario (main text) is slightly weaker than under directional selection for the optimum shifts we considered (up to two standard deviations of the trait distribution; Figure S10A).

#### S1.5 Gene-by-environment interactions (GxE)

##### Simulation Details

We simulated GxE using the same framework as the baseline scenario, except we drew the effect sizes of mutations from a multivariate normal distribution with three dimensions representing the ancestral burn-in population and its two descendant populations. The covariance matrix was determined according to the standard deviation of the effect size distribution and correlation of effect sizes provided (either 0.9 or 0.95). Phenotypes and thus fitness were calculated using the effect size assigned for the particular population in which they were evaluated. This means that the environments of the descendant populations changed at their time of divergence from the ancestral population, causing the effect sizes in each descendant population to be perturbed away from the ancestral effect sizes by a small normal deviate. These environmental changes do not generate a difference in the mean environment nor the optimum phenotype. The effect size correlations we chose are higher than most estimates, though GxG and differences in LD likely play a role as well (Liu *et al*., 2015; Galinsky *et al*., 2019; Lam *et al*., 2019; Veturi *et al*., 2019).

##### Accuracy of polygenic score predictions

Gene by environment interactions (GxE) combined with environmental differences between the GWAS sample and unrepresented population result in lower prediction accuracies in the unrepresented population, relative to the baseline scenario we discuss in the main text (Figure S10). Two factors contribute to why we find lower prediction accuracies under GxE. First, the effect sizes estimated by the GWAS in population A are only partially correlated with the true effects in population B. Thus, noise is introduced to the predictions when using the effect sizes from A for polygenic prediction in B. In Appendix A.2.1 we describe how we modify the equation for the reduction in prediction accuracy when the effect sizes between the GWAS sample and the prediction population are only partially correlated. Second, the slight changes in effect sizes, due to environmental changes experienced by each descendant population after they shared a common ancestor, transiently inflate the variance in a population relative to equilibrium conditions, leading to a faster removal of ancestral polymorphisms and thus fewer shared polymorphisms between the population pair. Specifically, at equilibrium, most common alleles have small effects. After an environmental change, many of these previously near-zero effect alleles tend to shift toward larger effect sizes (Figure S11). The accumulation of these minor shifts at many sites leads to an increase in the total variance within the descendant populations, and these changes are not correlated between them, so prediction accuracy is reduced.

##### Difference in polygenic means among populations

Under GxE, we also find higher chances of false signals of adaptation, even though the optimum never changes and is shared among populations. The chances of these false signals decay with time since the common ancestor of the population pair (Figure S13).

#### S1.6 Pleiotropy

##### Simulation Details

We simulated pleiotropy using the same framework as the baseline scenario. A mutation’s effect on each trait was independently drawn from the same normal distribution. An individual’s fitness was calculated using their Euclidean distance from the optimum, based on their distance in each phenotypic dimension. For this extension we simulated *n* = 5 and *n* = 10 traits (the baseline scenario is *n* = 1). The selection coefficient that an allele experiences is approximately

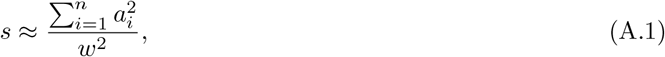

where *a*_*i*_ denotes the allele’s effect on trait *i* (Simons *et al*., 2018). Thus when we sample the effect of an allele on each trait independently and use the same distribution, with more pleiotropy there would always be stronger selection, where

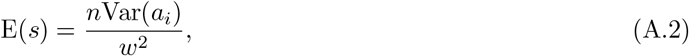

such that as *n* increases there would be a decrease in the amount of shared additive genetic variance, and thus portability, between populations. However, we were interested in investigating how the distribution of an allele’s effects on multiple traits impacted the amount of shared additive genetic variance between populations, in isolation from the effects of overall stronger selection on alleles. Therefore, we standardized the fitness function so that the average selection coefficient would be the same across pleiotropy levels (*n*). To achieve the same expectation across simulations, we multiplied *w*^2^ by *n* in the fitness function. While the average selection coefficient is the same for a particular *w* across *n*, its distribution differs according to the number of traits influenced,

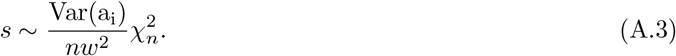

##### Accuracy of polygenic score predictions

Under this model of pleiotropy, we find less of a reduction in the prediction accuracy of ascertained variation compared to our baseline scenario. This is because the effect of the mutation on the trait of interest becomes more weakly correlated with its total selection coefficient. Specifically, common alleles can have larger effects on the trait of interest but by chance experience weaker selection due to small effects on the other selected traits. Therefore, the initial variance explained by a large effect polymorphism will be higher in the ancestor and there will be less of a per-generation reduction of variance at this locus. Together, more large effect and common alleles will be shared between populations, increasing the amount of additive genetic variance that polymorphisms ascertained in one population can explain in another. Thus pleiotropy, for a constant average strength of selection, reduces the rate at which prediction accuracy declines.

##### Difference in polygenic means among populations

Under pleiotropy, we see similar, though slightly lower, chances for false signals of adaptation as under the baseline stabilizing selection scenario (Figure S14).

**Figure S10:**
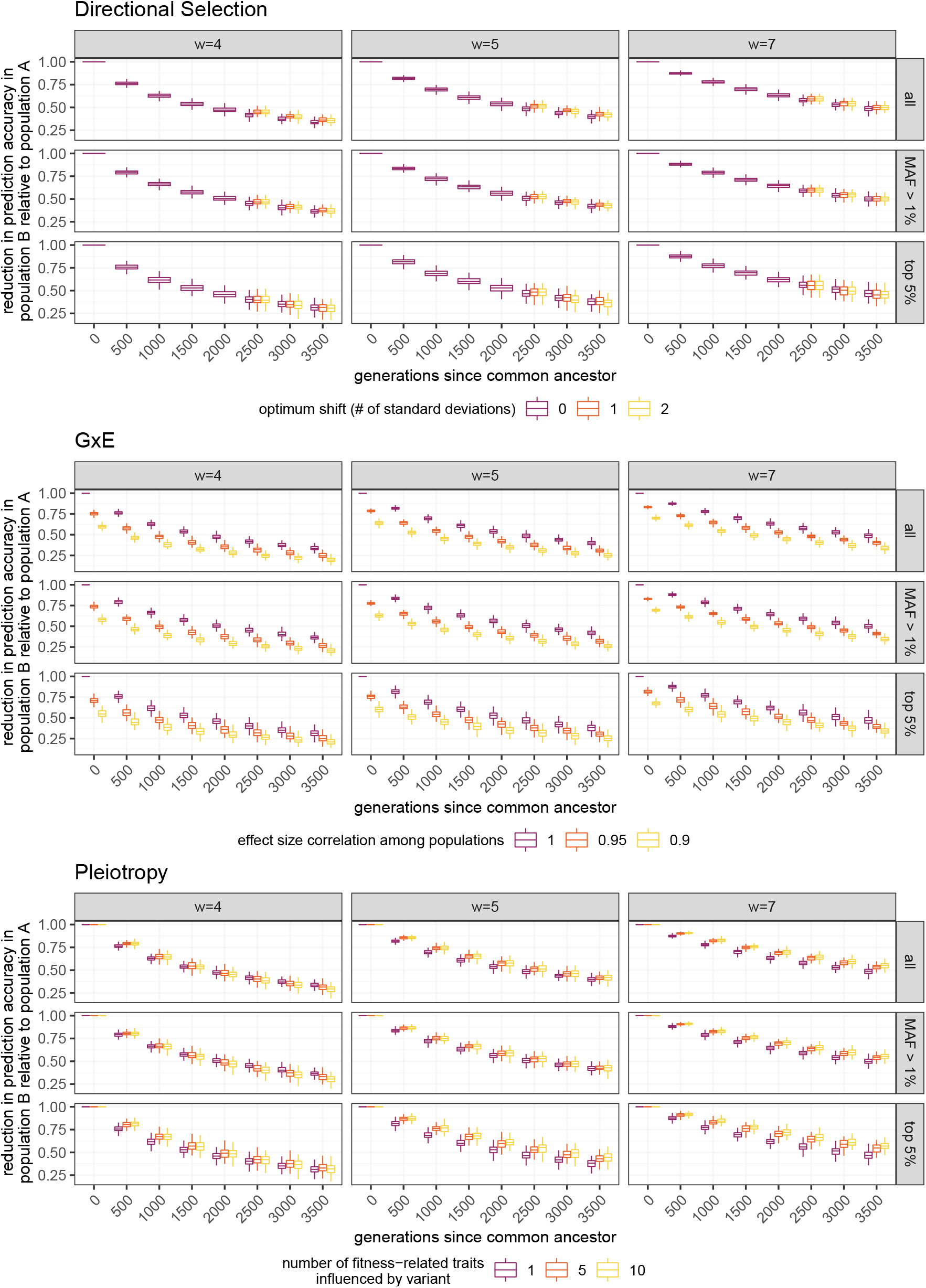
Comparison of the baseline stabilizing scenario and each extension (labelled by title) for the reduction in prediction accuracy in population B. Boxplots show the distribution of observations from 100 simulations, and are colored by the parameter values that vary in a particular extension. Results from the baseline scenario are always represented in purple (the first parameter value in each legend). Each column represents a different width of the fitness peak, and each row represents the set of polymorphisms ascertained in population A (*top*: all, *bottom*: top 5% based on variance contributed).

**Figure S11:**
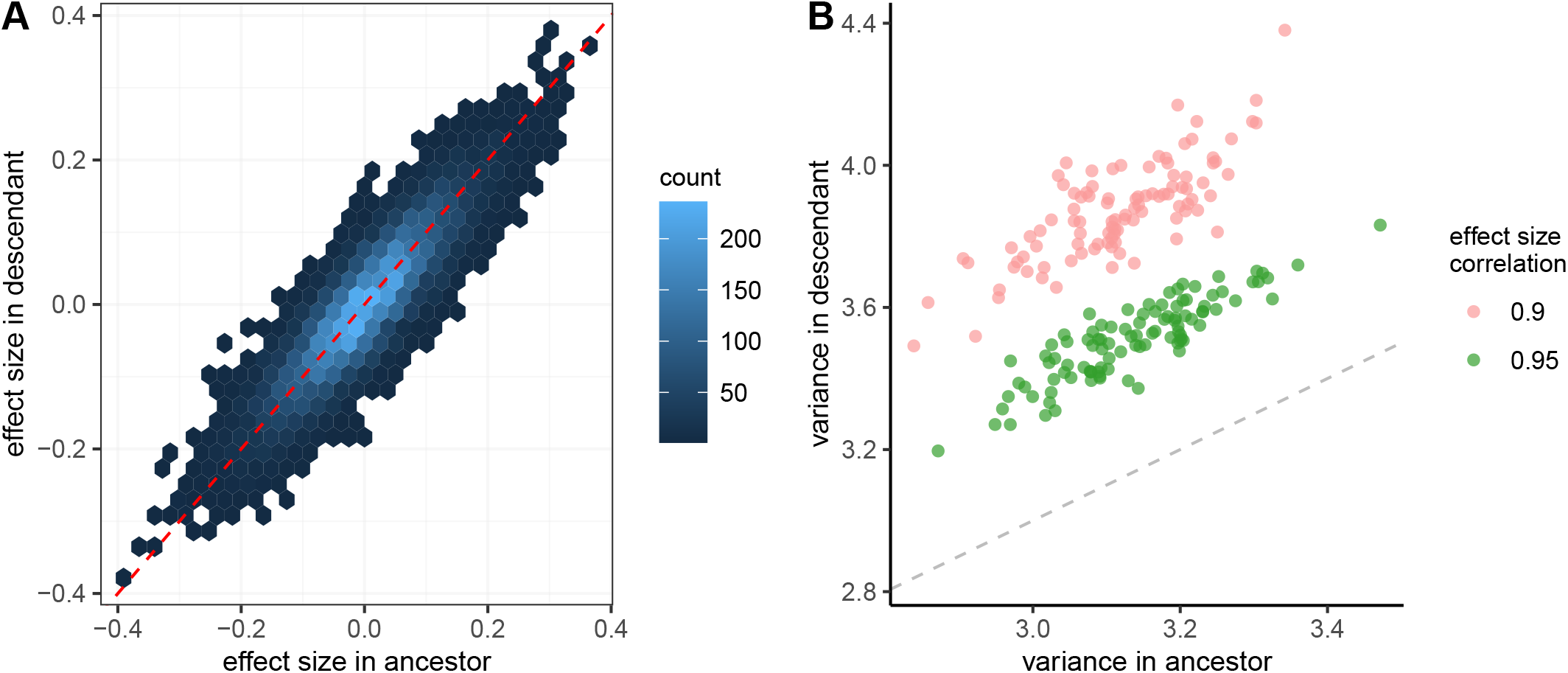
Shift of effect sizes and within-population variances when there’s GxE and populations A and B begin to experience different environments from their common ancestor at the time of divergence. **A)** Changes in effect sizes from the ancestral population (x-axis) to one of its descendant populations (y-axis) for each variance-contributing polymorphism. Colors represent the number of sites in a particular bin (outlined by hexagon). Results are shown for a single simulation with *w* = 5 and effect size correlation of 0.9. **B)** At the time of divergence/environmental shift, the variance in the descendant population (y-axis) increases relative to the variance in the ancestral population (x-axis) due to the shift in effect sizes. Each point represents a single simulation. Dashed lines in A and B represent lines of equality.

**Figure S12:**
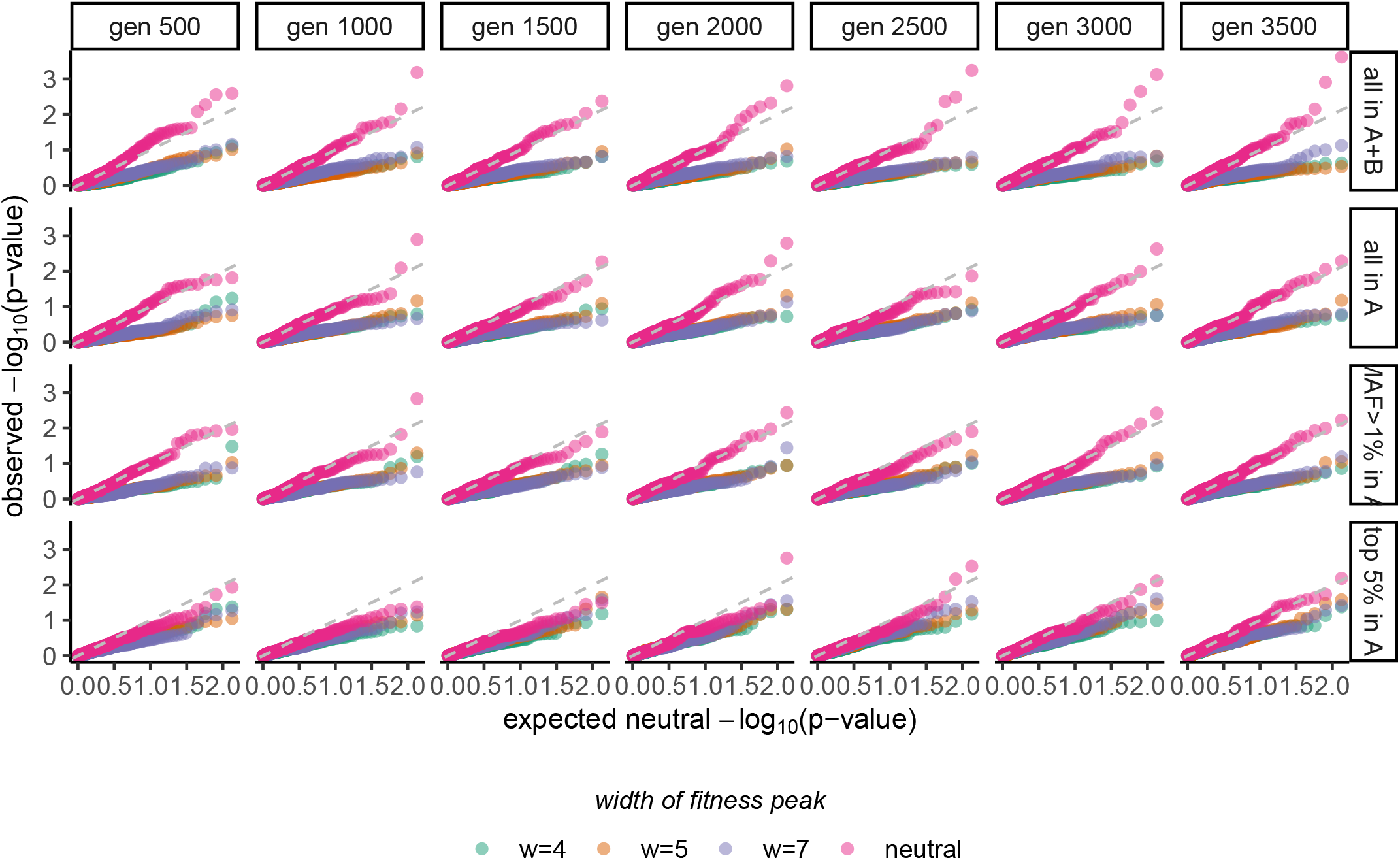
Quantile-quantile plot of observed p-values for *Q*_*X*_ under the **baseline** case against expected p-values under neutrality (uniform distribution). The dashed line shows equality; points that lie above this line indicate that the observed distribution has a higher density of low p-values than the neutral distribution, and points that lie below it indicate the opposite. Columns show results from every 500 generations of divergence between populations A and B and rows show results from different sets of ascertained polymorphims (*top:* all polymorphisms from both populations, *top-middle:* all polymorphisms in A, *bottom-middle:* polymorphisms with MAF*>*1% in A, *bottom:* top 5% of variance-contributing polymorphisms in A).

**Figure S13:**
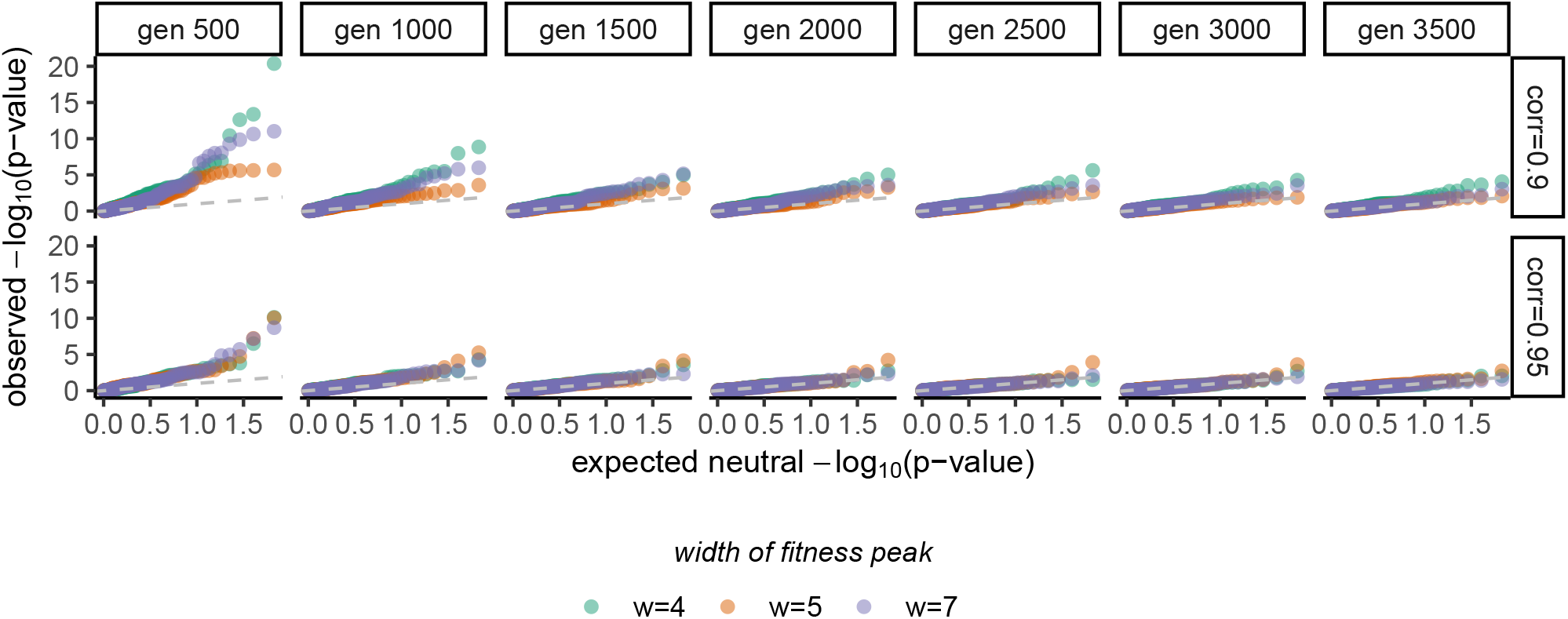
Quantile-quantile plot of observed p-values for *Q*_*X*_ under **GxE** against expected p-values under neutrality (uniform distribution). The dashed line shows equality; points that lie above this line indicate that the observed distribution has a higher density of low p-values than the neutral distribution, and points that lie below it indicate the opposite. Columns show results from every 500 generations of divergence between populations A and B and rows show results from each simulated correlation of effect sizes among populations.

**Figure S14:**
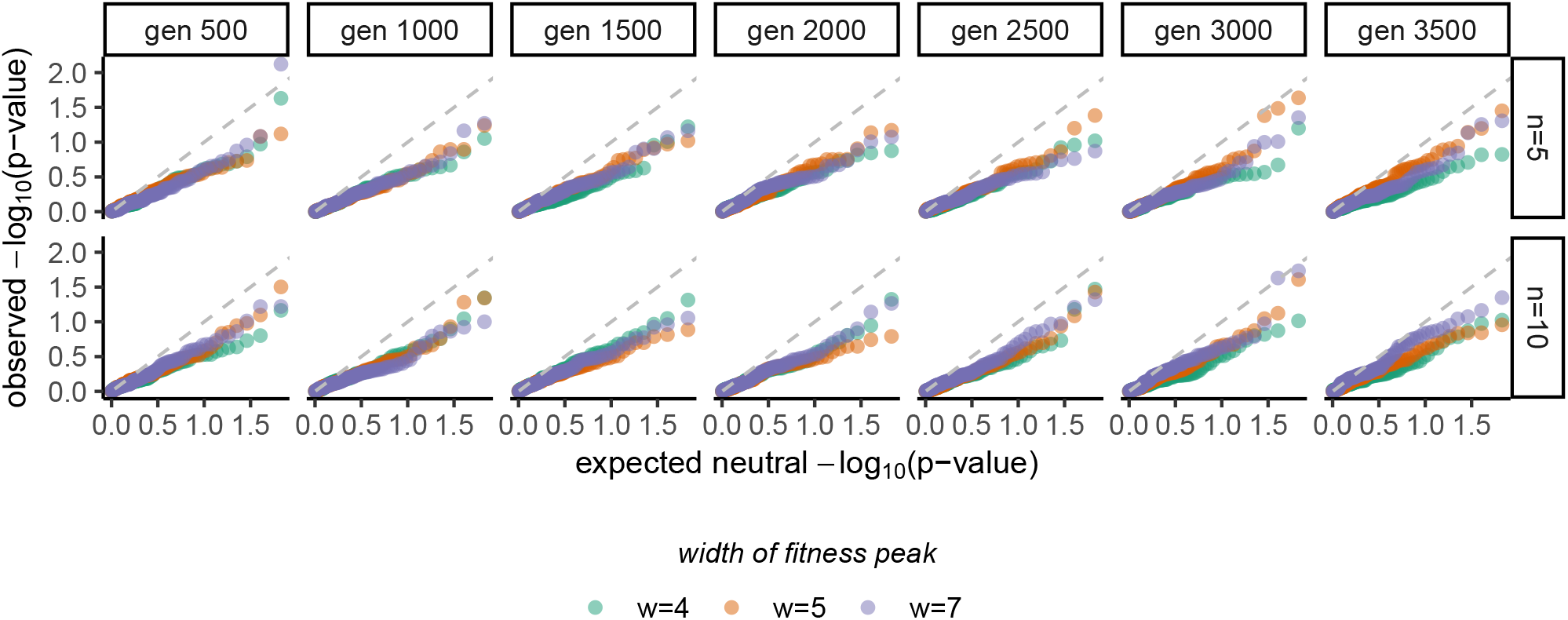
Quantile-quantile plot of observed p-values for *Q*_*X*_ under **pleiotropy** against expected p-values under neutrality (uniform distribution). The dashed line shows equality; points that lie above this line indicate that the observed distribution has a higher density of low p-values than the neutral distribution, and points that lie below it indicate the opposite. Columns show results from every 500 generations of divergence between populations A and B and rows show results from each simulated number of fitness-related traits (*n*) influenced by a mutation.

**Figure S15:**
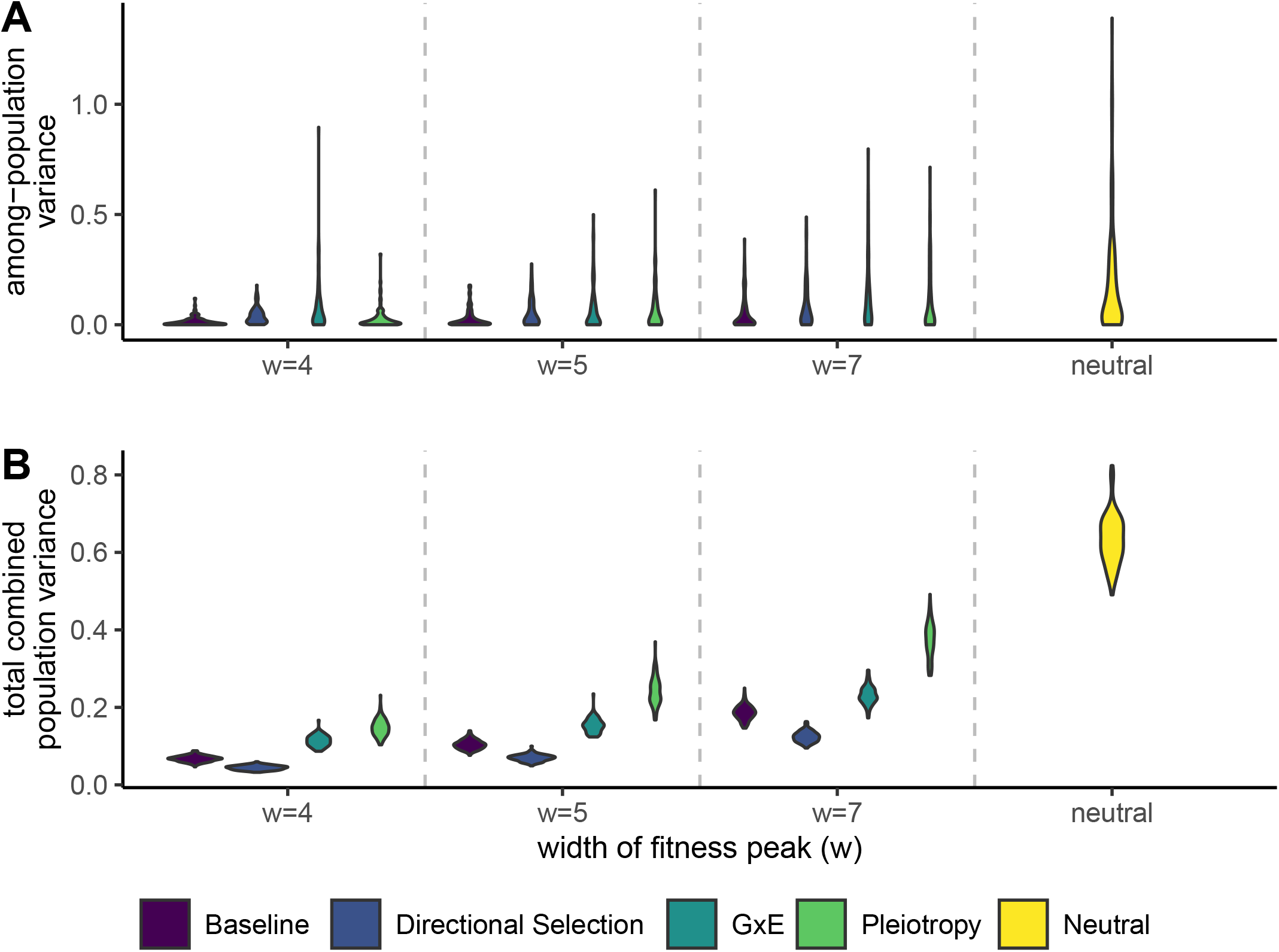
Components of *Q*_*ST*_ when estimated from the top 1% of variance-contributing sites in population A. Results for GxE are shown for the effect size correlation 0.9 and results for directional selection are shown for an optimum shift of two phenotypic standard deviations. **A)** The estimated mean difference in polygenic scores. The larger this value the larger the among-population variance, which is the numerator of *Q*_*ST*_. **B)** The total variance of the populations combined, which is the denominator of *Q*_*ST*_.

## References

Adhikari, K., J. Mendoza-Revilla, A. Sohail, M. Fuentes-Guajardo, J. Lampert, et al., 2019 A GWAS in Latin Americans highlights the convergent evolution of lighter skin pigmentation in Eurasia. Nature Communications 10.

Aris-Brosou, S., 2019 Direct Evidence of an Increasing Mutational Load in Humans. Molecular Biology and Evolution 36: 2823–2829.

Barbujani, G., A. Magagni, E. Minch, and L. L. Cavalli-Sforza, 1997 An apportionment of human dna diversity. Proceedings of the National Academy of Sciences 94: 4516–4519.

Barton, N. H. and P. D. Keightley, 2002 Understanding quantitative genetic variation. Nature Reviews Genetics 3: 11–21.

Bentley, A. R., Y. J. Sung, M. R. Brown, T. W. Winkler, A. T. Kraja, et al., 2019 Multi-ancestry genome-wide gene–smoking interaction study of 387,272 individuals identifies new loci associated with serum lipids. Nature Genetics 51: 636–648.

Berens, A. J., T. L. Cooper, and J. Lachance, 2017 The genomic health of ancient hominins. Human Biology 89: 7–19.

Berg, J. J. and G. Coop, 2014 A Population Genetic Signal of Polygenic Adaptation. PLoS Genetics 10: e1004412.

Berg, J. J., A. Harpak, N. Sinnott-Armstrong, A. M. Joergensen, H. Mostafavi, et al., 2019 Reduced signal for polygenic adaptation of height in UK Biobank. eLife 8: e39725.

Bergström, A., S. A. McCarthy, R. Hui, M. A. Almarri, Q. Ayub, et al., 2020 Insights into human genetic variation and population history from 929 diverse genomes. Science 367: eaay5012.

Bitarello, B. D. and I. Mathieson, 2020 Polygenic scores for height in admixed populations. G3: Genes, Genomes, Genetics 10: 4027–4036.

Boyle, E. A., Y. I. Li, and J. K. Pritchard, 2017 An Expanded View of Complex Traits: From Polygenic to Omnigenic. Cell 169: 1177–1186.

Brown, B. C., A. Genetic, E. Network, T. Diabetes, C. J. Ye, et al., 2016 Transethnic Genetic-Correlation Estimates from Summary Statistics. The American Journal of Human Genetics 99: 76–88.

Bulmer, M., 1971 The Effect of Selection on Genetic Variability. The American Naturalist 105: 201–211.

Bumpus, H., 1899 The Elimination of the Unfit as Illustrated by the Introduced Sparrow, Passer Domesticus: (a Fourth Contribution to the Study of Variation). Biological lectures delivered at the Marine Biological Laboratory of Wood’s Hole, Gin.

Carlson, M. O., D. P. Rice, J. J. Berg, and M. Steinrücken, 2021 Polygenic score accuracy in ancient samples: quantifying the effects of allelic turnover. bioRxiv.

Cavazos, T. B. and J. S. Witte, 2021 Inclusion of variants discovered from diverse populations improves polygenic risk score transferability. Human Genetics and Genomics Advances 2: 100017.

Chakraborty, R., 1990 Quantitative Traits in Relation to Population Structure: Why and How Are They Used and What Do They Imply? Human Biology 62: 147–162.

Chakraborty, R. and M. Nei, 1982 Genetic differentiation of quantitative characters between populations or species I: Mutation and random genetic drift. Genetical Research 39: 303–314.

Cohan, F. M., 1984 Can uniform selection retard random genetic divergence between isolated conspecific populations? Evolution 38: 495–504.

Conrad, D. F., M. Jakobsson, G. Coop, X. Wen, J. D. Wall, et al., 2006 A worldwide survey of haplotype variation and linkage disequilibrium in the human genome. Nature Genetics 38: 1251–1260.

Conti, D. V., B. F. Darst, L. C. Moss, E. J. Saunders, X. Sheng, et al., 2021 Trans-ancestry genome-wide association meta-analysis of prostate cancer identifies new susceptibility loci and informs genetic risk prediction. Nature Genetics 53: 65–75.

Coop, G., 2019 Reading tea leaves? polygenic scores and differences in traits among groups. arXiv preprint 1909.00892.

Coop, G., J. K. Pickrell, J. Novembre, S. Kudaravalli, J. Li, et al., 2009 The Role of Geography in Human Adaptation. PLoS Genetics 5: e1000500.

Coram, M. A., H. Fang, S. I. Candille, T. L. Assimes, and H. Tang, 2017 Leveraging Multi-ethnic Evidence for Risk Assessment of Quantitative Traits in Minority Populations. The American Journal of Human Genetics 101: 218–226.

Cox, S. L., H. Moots, J. T. Stock, A. Shbat, B. D. Bitarello, et al., 2021 Predicting skeletal stature using ancient DNA. bioRxiv.

Cox, S. L., C. B. Ruff, R. M. Maier, and I. Mathieson, 2019 Genetic contributions to variation in human stature in prehistoric Europe. Proceedings of the National Academy of Sciences 116: 201910606.

Curtis, D., 2018 Polygenic risk score for schizophrenia is more strongly associated with ancestry than with schizophrenia. Psychiatric Genetics 28: 85–89.

de Villemereuil, P., A. Charmantier, D. Arlt, P. Bize, P. Brekke, et al., 2020 Fluctuating optimum and temporally variable selection on breeding date in birds and mammals. Proceedings of the National Academy of Sciences 117: 31969–31978.

Duncan, L., H. Shen, B. Gelaye, J. Meijsen, K. Ressler, et al., 2019 Analysis of polygenic risk score usage and performance in diverse human populations. Nature Communications 10.

Durvasula, A. and K. E. Lohmueller, 2021 Negative selection on complex traits limits phenotype prediction accuracy between populations. The American Journal of Human Genetics 108: 620–631.

Edge, M. D. and N. A. Rosenberg, 2015 A general model of the relationship between the apportionment of human genetic diversity and the apportionment of human phenotypic diversity. Human Biology 87: 313–337.

Esteller-Cucala, P., I. Maceda, A. D. Børglum, D. Demontis, S. V. Faraone, et al., 2020 Genomic analysis of the natural history of attention-deficit/hyperactivity disorder using Neanderthal and ancient Homo sapiens samples. Scientific Reports 10.

Fan, S., M. E. Hansen, Y. Lo, and S. A. Tishkoff, 2016 Going global by adapting local: A review of recent human adaptation. Science 354: 54–59.

Feldman, M. W. and R. C. Lewontin, 1975 The Heritability Hang-up. Science 190: 1163–1168.

Galinsky, K. J., Y. A. Reshef, H. K. Finucane, P. R. Loh, N. Zaitlen, et al., 2019 Estimating cross-population genetic correlations of causal effect sizes. Genetic Epidemiology 43: 180–188.

Gingerich, P. D., 1983 Rates of evolution: Effects of time and temporal scaling. Science 222: 159–161.

Goldstein, D. B. and K. E. Holsinger, 1992 Maintenance of polygenic variation in spatially structured populations: roles for local mating and genetic redundancy. Evolution 46: 412–429.

Grinde, K. E., T. A. Thornton, K. H. K. Chan, and A. P. Reiner, 2019 Generalizing polygenic risk scores from Europeans to Hispanics / Latinos. Genetic Epidemiology 43: 50–62.

Haller, B. C. and P. W. Messer, 2019 SLiM 3: Forward Genetic Simulations Beyond the Wright-Fisher Model. Molecular Biology and Evolution 36: 632–637.

Harpak, A. and M. Przeworski, 2021 The evolution of group differences in changing environments. PLoS Biology 19: e3001072.

Haworth, S., R. Mitchell, L. Corbin, K. H. Wade, T. Dudding, et al., 2019 Apparent latent structure within the UK Biobank sample has implications for epidemiological analysis. Nature Communications 10.

Hayward, L. K. and G. Sella, 2021 Polygenic adaptation after a sudden change in environment. bioRxiv.

Hernandez, R. D., J. L. Kelley, E. Elyashiv, S. C. Melton, A. Auton, et al., 2011 Classic Selective Sweeps Were Rare in Recent Human Evolution. Science 257: 920–924.

Hill, W. G., M. E. Goddard, and P. M. Visscher, 2008 Data and theory point to mainly additive genetic variance for complex traits. PLOS Genetics 4: 1–10.

Hill, W. G. and M. Kirkpatrick, 2010 What animal breeding has taught us about evolution. Annual Review of Ecology, Evolution, and Systematics 41: 1–19.

Horikoshi, M., T. Sofer, A. Mahajan, H. Kitajima, N. Franceschini, et al., 2017 Trans-ethnic meta-regression of genome-wide association studies accounting for ancestry increases power for discovery and improves fine-mapping resolution. Human Molecular Genetics 26: 3639–3650.

Houle, D., G. H. Bolstad, K. Van Der Linde, and T. F. Hansen, 2017 Mutation predicts 40 million years of fly wing evolution. Nature 548: 447–450.

Irving-Pease, E. K., R. Muktupavela, M. Dannemann, and F. Racimo, 2021 Quantitative Human Paleogenetics: what can ancient DNA tell us about complex trait evolution? Frontiers in Genetics 12: 703541.

Isshiki, M., Y. Watanabe, and J. Ohashi, 2021 Geographic variation in the polygenic score of height in Japan. Human Genetics 140: 1097–1108.

Jorde, L. B., W. S. Watkins, M. J. Bamshad, M. Dixon, C. Ricker, et al., 2000 The distribution of human genetic diversity: a comparison of mitochondrial, autosomal, and y-chromosome data. The American Journal of Human Genetics 66: 979–988.

Keightley, P. D. and W. G. Hill, 1988 Quantitative genetic variability maintained by mutation-stabilizing selection balance in finite populations. Genetical Research 52: 33–43.

Kerminen, S., A. R. Martin, J. Koskela, S. E. Ruotsalainen, A. S. Havulinna, et al., 2019 Geographic Variation and Bias in the Polygenic Scores of Complex Diseases and Traits in Finland. American Journal of Human Genetics 104: 1169–1181.

Kim, M. S., K. P. Patel, A. K. Teng, A. J. Berens, and J. Lachance, 2018 How genetic disease risks can be misestimated across global populations. Genome Biology 19.

Kingsolver, J. G., H. E. Hoekstra, J. M. Hoekstra, D. Berrigan, S. N. Vignieri, et al., 2001 The strength of phenotypic selection in natural populations. The American Naturalist 157: 245–261.

Kremer, A. and V. Le Corre, 2012 Decoupling of differentiation between traits and their underlying genes in response to divergent selection. Heredity 108: 375–385.

Lam, M., C.-y. Chen, Z. Li, A. R. Martin, and J. Bryois, 2019 Comparative genetic architectures of schizophrenia in East Asian and European populations. Nature Genetics 51.

Lande, R., 1976 Natural selection and random genetic drift in phenotypic evolution. Evolution 30: 314–334.

Lande, R., 1991 Isolation by distance in a quantitative trait. Genetics 128: 443–452.

Lande, R., 1992 Neutral Theory of Quantitative Genetic Variance in an Island Model with Local Extinction and Colonization 46: 381–389.

Lande, R. and S. J. Arnold, 1983 The measurement of selection on correlated characters. Evolution 37: 1210–1226.

Latta, R. G., 1998 Differentiation of allelic frequencies at quantitative trait loci affecting locally adaptive traits. American Naturalist 151: 283–292.

Le Corre, V. and A. Kremer, 2003 Genetic Variability at Neutral Markers, Quantitative Trait Loci and Trait. Genetics 1219: 1205–1219.

Lewontin, R. C., 1972 The Apportionment of Human Diversity. In Evolutionary Biology, edited by T. Dobzhansky, M. K. Hecht, and W. C. Steere, chapter 14, pp. 381–398, Appleton-Century-Crofts, New York.

Lewontin, R. C., 1974 The analysis of variance and the analysis of causes. American Journal of Human Genetics 26: 400–411.

Lewontin, R. C. and J. Krakauer, 1973 Distribution of gene frequency as a test of the theory of the selective neutrality of polymorphisms. Genetics 74: 175–195.

Li, J. Z., D. M. Absher, H. Tang, A. M. Southwick, A. M. Casto, et al., 2008 Worldwide human relationships inferred from genome-wide patterns of variation. Science 319: 1100–1104.

Li, Y. R. and B. J. Keating, 2014 Trans-ethnic genome-wide association studies: advantages and challenges of mapping in diverse populations 6.

Liu, J. Z., S. Van Sommeren, H. Huang, S. C. Ng, R. Alberts, et al., 2015 Association analyses identify 38 susceptibility loci for inflammatory bowel disease and highlight shared genetic risk across populations. Nature Genetics 47: 979–986.

Loh, P. R., G. Bhatia, A. Gusev, H. K. Finucane, B. K. Bulik-Sullivan, et al., 2015 Contrasting genetic architectures of schizophrenia and other complex diseases using fast variance-components analysis. Nature Genetics 47: 1385–1392.

Macarthur, J., E. Bowler, M. Cerezo, L. Gil, P. Hall, et al., 2017 The new NHGRI-EBI Catalog of published genome-wide association studies (GWAS Catalog). Nucelic Acids Research 45: 896–901.

Marciniak, S., C. M. Bergey, A. M. Silva, A. Haluszko, M. Furmanek, et al., 2021 An integrative skeletal and paleogenomic analysis of prehistoric stature variation suggests relatively reduced health for early European farmers. bioRxiv.

Márquez-Luna, C., 2017 Multiethnic polygenic risk scores improve risk prediction in diverse populations. Genetic Epidemiology 41: 811–823.

Martin, A. R., C. R. Gignoux, R. K. Walters, G. L. Wojcik, B. M. Neale, et al., 2017a Human Demographic History Impacts Genetic Risk Prediction across Diverse Populations. The American Journal of Human Genetics 100: 635–649.

Martin, A. R., M. Kanai, Y. Kamatani, Y. Okada, B. M. Neale, et al., 2019 Clinical use of current polygenic risk scores may exacerbate health disparities. Nature Genetics 51: 584–591.

Martin, A. R., M. Lin, J. M. Granka, J. W. Myrick, X. Liu, et al., 2017b An Unexpectedly Complex Architecture for Skin Pigmentation in Africans. Cell 171: 1340–1353.

Martiniano, R., L. M. Cassidy, R. Ó ‘Maoldúin, R. McLaughlin, N. M. Silva, et al., 2017 The population genomics of archaeological transition in west Iberia: Investigation of ancient substructure using imputation and haplotype-based methods. PLoS Genetics 13: 1–24.

Maruyama, T. and M. Kimura, 1975 Moments for Sum of an Arbitrary Function of Gene Frequency along a Stochastic Path of Gene Frequency Change. Proceedings of the National Academy of Sciences 72: 1602–1604.

Mathieson, I., 2021 The omnigenic model and polygenic prediction of complex traits. The American Journal of Human Genetics 108: 1–6.

Mathieson, I., I. Lazaridis, N. Rohland, S. Mallick, N. Patterson, et al., 2015 Genome-wide patterns of selection in 230 ancient Eurasians. Nature 528: 499–503.

Mostafavi, H., A. Harpak, I. Agarwal, D. Conley, J. K. Pritchard, et al., 2020 Variable prediction accuracy of polygenic scores within an ancestry group. eLife 9: e48376.

Narain, P. and R. Chakraborty, 1987 Genetic differentiation of quantitative characters between populations or species II: Optimal selection in infinite populations. Heredity 59: 199–212.

Novembre, J. and N. H. Barton, 2018 Tread lightly interpreting polygenic tests of selection. Genetics 208: 1351–1355.

Patel, R. A., S. A. Musharoff, J. P. Spence, H. Pimentel, C. Tcheandjieu, et al., 2021 Effect sizes of causal variants for gene expression and complex traits differ between populations. bioRxiv.

Popejoy, A. B. and S. M. Fullerton, 2016 Genomics is failing on diversity. Nature 538: 161–164.

Privé, F., H. Aschard, S. Carmi, L. Folkersen, C. Hoggart, et al., 2022 Portability of 245 polygenic scores when derived from the uk biobank and applied to 9 ancestry groups from the same cohort. The American Journal of Human Genetics 109: 12–23.

Prout, T. and J. S. F. Barker, 1993 F Statistics in Drosophila buzzatii: Selection, Population Size and Inbreeding. Genetics 134: 369–375.

Refoyo-Martínez, A., S. Liu, A. M. Jørgensen, X. Jin, A. Albrechtsen, et al., 2020 How robust are cross-population signatures of polygenic adaptation in humans? bioRxiv pp. 1–66.

Relethford, J. H. and F. C. Lees, 1982 The use of quantitative traits in the study of human population structure. American Journal of Physical Anthropology 25: 113–132.

Robertson, A., 1956 The effect of selection against extreme deviants based on deviation or on homozygosis. Journal of Genetics 54: 236–248.

Rogers, A. R. and H. C. Harpending, 1983 Population structure and quantitative characters. Genetics 105: 985–1002.

Rose, G., 2001 Sick individuals and sick populations. International Journal of Epidemiology 30: 396–427–432.

Rosenberg, N. A., M. D. Edge, J. K. Pritchard, and M. W. Feldman, 2019 Interpreting polygenic scores, polygenic adaptation, and human phenotypic differences. Evolution, medicine, and public health 2019: 26–34.

Rosenberg, N. A., J. K. Pritchard, J. L. Weber, H. M. Cann, K. K. Kidd, et al., 2002 Genetic structure of human populations. Science 298: 2381–2385.

Sakaue, S., M. Kanai, Y. Tanigawa, J. Karjalainen, M. Kurki, et al., 2020 A global atlas of genetic associations of 220 deep phenotypes. medRxiv pp. 1–52.

Sanjak, J. S., J. Sidorenko, M. R. Robinson, K. R. Thornton, and P. M. Visscher, 2018 Evidence of directional and stabilizing selection in contemporary humans. Proceedings of the National Academy of Sciences 115: 151–156.

Sawyer, S. A. and D. L. Hartl, 1992 Population Genetics of Polymorphism and Divergence. Genetics 132: 1161–1176.

Scutari, M., I. Mackay, and D. Balding, 2016 Using genetic distance to infer the accuracy of genomic prediction. PLOS Genetics 12: 1–19.

Sella, G. and N. H. Barton, 2019 Thinking about the evolution of complex traits in the era of genome-wide association studies. Annual Review of Genomics and Human Genetics 20: 461–493.

Shi, H., G. Kichaev, and B. Pasaniuc, 2016 Contrasting the Genetic Architecture of 30 Complex Traits from Summary Association Data. American Journal of Human Genetics 99: 139–153.

Simons, Y. B., K. Bullaughey, R. R. Hudson, and G. Sella, 2018 A population genetic interpretation of GWAS findings for human quantitative traits. PLoS Biology 16.

Simonti, C. N. and J. Lachance, 2021 Ancient DNA reveals that few GWAS loci have been strongly selected during recent human history. bioRxiv.

Sohail, M., R. M. Maier, A. Ganna, A. Bloemendal, A. R. Martin, et al., 2019 Polygenic adaptation on height is overestimated due to uncorrected stratification in genome-wide association studies. eLife 8: e39702.

Spitze, K., 1993 Population structure in Daphnia obtusa: Quantitative genetic and allozymic variation. Genetics 135: 367–374.

Trochet, H. and J. Hussin, 2020 Fine-scale population structure confounds genetic risk scores in the ascertainment population. bioRxiv.

Turchin, M. C., C. W. Chiang, C. D. Palmer, S. Sankararaman, D. Reich, et al., 2012 Evidence of widespread selection on standing variation in europe at height-associated snps. Nature Genetics 44: 1015–1019.

Turelli, M., 1984 Heritable genetic variation via mutation-selection balance: Lerch’s zeta meets the abdominal bristle. Theoretical Population Biology 25: 138–193.

Turelli, M. and N. Barton, 1990 Dynamics of polygenic characters under selection. Theoretical Population Biology 38: 1–57.

Turelli, M. and N. H. Barton, 1994 Genetic and statistical analyses of strong selection on polygenic traits: what, me normal? Genetics 138: 913–941.

Veturi, Y., N. Yi, W. Huang, and A. I. Vazquez, 2019 Modeling Heterogeneity in the Genetic Architecture of Ethnically Diverse Groups Using Random Effect Interaction Models. Genetics 211: 1395–1407.

Vilhjálmsson, B. J., J. Yang, H. K. Finucane, A. Gusev, S. Ripke, et al., 2015 Modeling Linkage Disequilibrium Increases Accuracy of Polygenic Risk Scores. American Journal of Human Genetics 97: 576–592.

Wang, Y., J. Guo, G. Ni, J. Yang, P. M. Visscher, et al., 2020 Theoretical and empirical quantification of the accuracy of polygenic scores in ancestry divergent populations. Nature Communications 11: 1–9.

Weber, K. E., 1990 Selection on wing allometry in Drosophila melanogaster. Genetics 126: 975–989.

Weissbrod, O., M. Kanai, H. Shi, S. Gazal, W. Peyrot, et al., 2021 Leveraging fine-mapping and non-European training data to improve trans-ethnic polygenic risk scores. medRxiv.

Whitlock, M. C., 1999 Neutral additive genetic variance in a metapopulation. Genetics Research 74: 215–221.

Wojcik, G. L., M. Graff, K. K. Nishimura, R. Tao, J. Haessler, et al., 2019 Genetic analyses of diverse populations improves discovery for complex traits. Nature 570: 514–518.

Wood, A. R., T. Esko, J. Yang, S. Vedantam, T. H. Pers, et al., 2014 Wood, Andrew R Esko, Tonu Yang, Jian Vedantam S, Pers TH, Gustafsson S, et al. Nature Genetics 46.

Wright, S., 1935 Evolution in populations in approximate equilibrium. Journal of Genetics 30: 257–266.

Wright, S., 1951 The Genetical Structure of Populations. Annals of Eugenics 15: 323–354.

Yang, J., A. Bakshi, Z. Zhu, G. Hemani, A. A. E. Vinkhuyzen, et al., 2015 Genetic variance estimation with imputed variants finds negligible missing heritability for human height and body mass index. Nature Genetics 47.

Yang, J., B. Benyamin, B. P. Mcevoy, S. Gordon, A. K. Henders, et al., 2010 Common SNPs explain a large proportion of the heritability for human height. Nature Genetics 42.

